# Using *de novo* assembly to identify structural variation of complex immune system gene regions

**DOI:** 10.1101/2021.02.03.429586

**Authors:** Jia-Yuan Zhang, Hannah Roberts, David S. C. Flores, Antony J. Cutler, Andrew C. Brown, Justin P. Whalley, Olga Mielczarek, David Buck, Helen Lockstone, Barbara Xella, Karen Oliver, Craig Corton, Emma Betteridge, Rachael Bashford-Rogers, Julian C. Knight, John A. Todd, Gavin Band

**Author notes:** Corresponding authors (JYZ); (GB). Senior authors (Julian C. Knight, John A. Todd, Gavin Band).

## Abstract

Driven by the necessity to survive environmental pathogens, the human immune system has evolved exceptional diversity and plasticity, to which several factors contribute including inheritable structural polymorphism of the underlying genes. Characterizing this variation is challenging due to the complexity of these loci, which contain extensive regions of paralogy, segmental duplication and high copy-number repeats, but recent progress in long-read sequencing and optical mapping techniques suggests this problem may now be tractable. Here we assess this by using long-read sequencing platforms from PacBio and Oxford Nanopore, supplemented with short-read sequencing and Bionano optical mapping, to sequence DNA extracted from CD14^+^ monocytes and peripheral blood mononuclear cells from a single European individual identified as HV31. We use this data to build a *de novo* assembly of eight genomic regions encoding four key components of the immune system, namely the human leukocyte antigen, immunoglobulins, T cell receptors, and killer-cell immunoglobulin-like receptors. Validation of our assembly using k-mer based and alignment approaches suggests that it has high accuracy, with estimated base-level error rates below 1 in 10 kb, although we identify a small number of remaining structural errors. We use the assembly to identify heterozygous and homozygous structural variation in comparison to GRCh38. Despite analyzing only a single individual, we find multiple large structural variants affecting core genes at all three immunoglobulin regions and at two of the three T cell receptor regions. Several of these variants are not accurately callable using current algorithms, implying that further methodological improvements are needed. Our results demonstrate that assessing haplotype variation in these regions is possible given sufficiently accurate long-read and associated data; application of these methods to larger samples would provide a broader catalogue of germline structural variation at these loci, an important step toward making these regions accessible to large-scale genetic association studies.

## Introduction

The capability of the human immune system to respond to environmental pathogens results from its substantial diversity and variability, both among individuals within a population and among cells within a single host. Key components of the innate and adaptive immune system, including the human leukocyte antigen (HLA), immunoglobulins (IG), T cell receptors (TCR) and killer-cell immunoglobulin-like receptors (KIR), have evolved exceptional complexity in their genomic loci, featuring numerous highly similar genes interspersed with pseudogenes and repetitive elements. Variation in genes encoding some of these components have well-established associations with infectious, immune-mediated, and other disease traits. The major histocompatibility complex (MHC) encoding HLA is so far the best-studied example, with hundreds of associations now known across multiple classes of disease [1,2] including infections [3,4]. In some cases the underlying functional mechanisms have also been identified [5,6]. However, despite the clearly important role of immunoglobulins (IG), TCR and KIR [7– 9], the underlying complexity of these genomic regions has so far prevented a full analysis of their contribution to human disease.

Three challenges must be overcome to make these regions accessible to future studies. First, key aspects of adaptive immunity are driven by somatic recombination and hypermutation of TCR and IG genes in immune cells. Consequently, DNA from non-recombining cell populations is needed to access germline genetic variation in these regions; these are not targeted in current surveys of haplotype variation based on lymphoblastoid cell lines or whole blood [10–12]. Second, extensive paralogy makes these regions intractable to short-read sequencing approaches [13], although analyses based on known immunogenetic sequences can be achieved [14]. Approaches using more costly long-read sequencing must therefore be employed [15], though even these methods do not currently solve the most complex regions [16]. Third, even if these technical challenges can be dealt with, the high diversity observed at these regions presents further difficulties for methods that identify, catalogue, and genotype the underlying variation. Solving these challenges would in principle enable the development of large haplotype variation reference panels at these loci, complementing existing immunogenetic variation databases [17] and opening them to analysis in large disease association studies.

Motivated by these challenges, here we utilize genomic data from a single individual (HV31) to assemble eight regions that encode key components of the human adaptive and innate immune response. To achieve this, we use DNA extracted from CD14^+^ monocytes, which do not undergo systematic somatic recombination. We develop a pipeline that exploits PacBio circular consensus sequencing (CCS), Bionano optical mapping, and short-read sequencing data to produce high-quality *de novo* assemblies of these regions. We then use additional long-read and short-read datasets to assess assembly accuracy and to call heterozygous variations using computational approaches based on read alignment and the copy number distribution of short k-mers (i.e. short DNA fragments of fixed length k). We find that HV31 carries substantial structural differences between haplotypes and in comparison to the GRCh38 reference sequence, including multiple large variants that affect core immune system genes but are not accurately called by current methods, and we investigate several of these in detail. Lastly, we analyze four gaps in the GRCh38 reference sequence at the immunoglobulin κ and T cell receptor γ regions, that are fully or partially filled in our assembly.

## Results

### Immune system loci display a spectrum of complexity in the human reference sequence

We focused on eight genome regions that encode components of the human immune system, namely those encoding the HLA, immunoglobulins (IGH, IGL, IGK), T cell receptors (TRA, TRB, TRD, TRG), and the killer-cell immunoglobulin-like receptors (KIR) (Table 1). Regions were defined based on NCBI RefSeq locus definitions [18] (except HLA and KIR which were based on previously published gene ranges [19,20]), plus an additional 1Mb flanking sequence added to both sides (see Methods). In the IGK region we additionally expanded the range to include a ∼1 Mb heterochromatin gap present in GRCh38. The expanded regions range from 2-6 Mb in length and vary considerably in terms of repetitive structure and haplotype diversity (Table 1). We noted the least reference sequence complexity in the T cell receptor α, d and γ regions (which contain < 2% repeat sequence and no listed alternate haplotypes), but greater complexity in other regions. In particular, the regions encoding immunoglobulin subunits contain the highest levels of duplication; previous analyses [21,22] have demonstrated significant structural diversity among known haplotypes in these regions. GRCh38 also contains dozens of alternative haplotype sequences at the HLA and KIR regions, and four gaps in the IGK and TRB regions (Table 1). Comparison to the earlier GRCh37 assembly and the presence of fix patches highlights that these regions are likely to be challenging to assemble.

**Table 1.** Overview of eight selected immune system loci in GRCh38.

| Name | Acronym | Coordinates and length | # core genes | % repetitive<sup>a</sup> | % SD<sup>b</sup> | # Gaps<sup>c</sup> | # Alternate haplotypes | % Novel to GRCh38<sup>d</sup> | # Fix patches<sup>e</sup> |
| --- | --- | --- | --- | --- | --- | --- | --- | --- | --- |
| Immuno-globulin heavy chain | IGH | chr14<br>104,586,437 - 107,043,718<br>(2.46Mb) | 164 | 6.8 | 31.1<br>(0.5) | 0 | 2 | 44.7 | 0 |
| Immuno-globulin $\kappa$ | IGK | chr2<br>87,857,361 - 9,1902,511<br>(4.05Mb) | 84 | 22.7 | 44.8<br>(22.7) | 3 | 2 | 31.5 | 1 |
| Immuno-globulin $\lambda$ | IGL | chr22<br>21,026,076 - 23,922,913<br>(2.90Mb) | 89 | 7.4 | 34.0<br>(15.3) | 0 | 3 | 47.4 | 0 |
| Human leukocyte antigen | HLA | chr6<br>28,602,238 - 34,409,896<br>(5.81Mb) | 39 | 2.7 | 6.5<br>(1.1) | 0 | 8 | 0 | 0 |
| T cell receptor $\alpha$ and $\delta$ | TRA | chr14<br>20,621,904 - 23,552,132<br>(2.93Mb) | 115 | 1.9 | 3.5 (0) | 0 | 0 | 37.6 | 0 |
| T cell receptor $\beta$ | TRB | chr7<br>141,299,011 - 143,813,287<br>(2.51Mb) | 78 | 5.3 | 19.5<br>(9.0) | 1 | 2 | 34.2 | 1 |
| T cell receptor $\gamma$ | TRG | chr7<br>37,240,024 - 39,368,055<br>(2.13Mb) | 22 | 1.3 | 3.1 (0) | 0 | 0 | 0.2 | 0 |
| Killer-cell immunoglobulin-like receptors | KIR | chr19<br>53,724,447 - 55,867,209<br>(2.14Mb) | 10 | 4.7 | 12.9<br>(0) | 0 | 50 | 47.4 | 0 |

<sup>a</sup> The proportion of repetitive DNA calculated as the proportion of 31-mers that are repeated at least once.
<sup>b</sup> The percentage of the region that is annotated as lying in a segmental duplication or (in brackets) a highly identical ( $\geq 95\%$ ) segmental duplication.
<sup>c</sup> The number of gaps (sequences of ‘N’ bases) in GRCh38.
<sup>d</sup> The percentage length of contigs that are new to GRCh38, i.e. not carried forward from GRCh37.
<sup>e</sup> The number of fix patches intersecting the locus in GRCh38 patch release 13.

### Assembling immune system regions with long-read, short-read and optical mapping data

We assessed whether the eight selected regions can be accurately assembled *de novo* using data from a single individual identified here as HV31. HV31 was recruited as a healthy volunteer and identified as having European ancestry. To facilitate accurate assembling of these complex regions, we generated data from multiple complementary platforms. Specifically, we performed PacBio Sequel II CCS (obtaining 12.3× genome coverage by ∼12 kb reads), MGI short-read sequencing (56.8×) and Bionano Saphyr Direct Label and Stain (DLS) optical mapping (152.7× coverage by imaged molecules). In addition, long-read and short-read sequencing data from PacBio continuous long read (CLR; 35×), Oxford Nanopore Technologies (ONT) PromethION (63×), 10x Genomics linked-reads (40.2×), Illumina Novaseq PCR-free (44.2×), MGI single-tube long fragment read (stLFR) (51.3×) and MGI CoolMPS (56.9×) platforms were also generated from the same blood sample (S1 Table). To minimize the impact of cell-specific events including V(D)J recombination and somatic hypermutation and enable accurate assembly of the germline genome, the data used for our assembly below were collected from CD14^+^ monocytes isolated from peripheral blood mononuclear cells (PBMCs) with antibody-conjugated beads (see Methods). As an exception, the Bionano imaging data was generated directly from PBMCs, as DNA from CD14^+^ monocytes failed to yield high-quality imaging data. The data generated in this study are further detailed in S1 Table, and will be made available through the European Genome-phenome Archive (EGA; see Data Availability).

To generate an accurate representation of the eight regions in the HV31 genome, we developed an assembly pipeline consisting of four stages (Fig 1A). This approach deals with heterozygosity by producing a consensus assembly of each region, and a list of heterozygous structural variants (SVs; Fig 1B and S2 Dataset) that jointly describe the HV31 genome. We describe the steps of this assembly process below.

**Fig 1.**
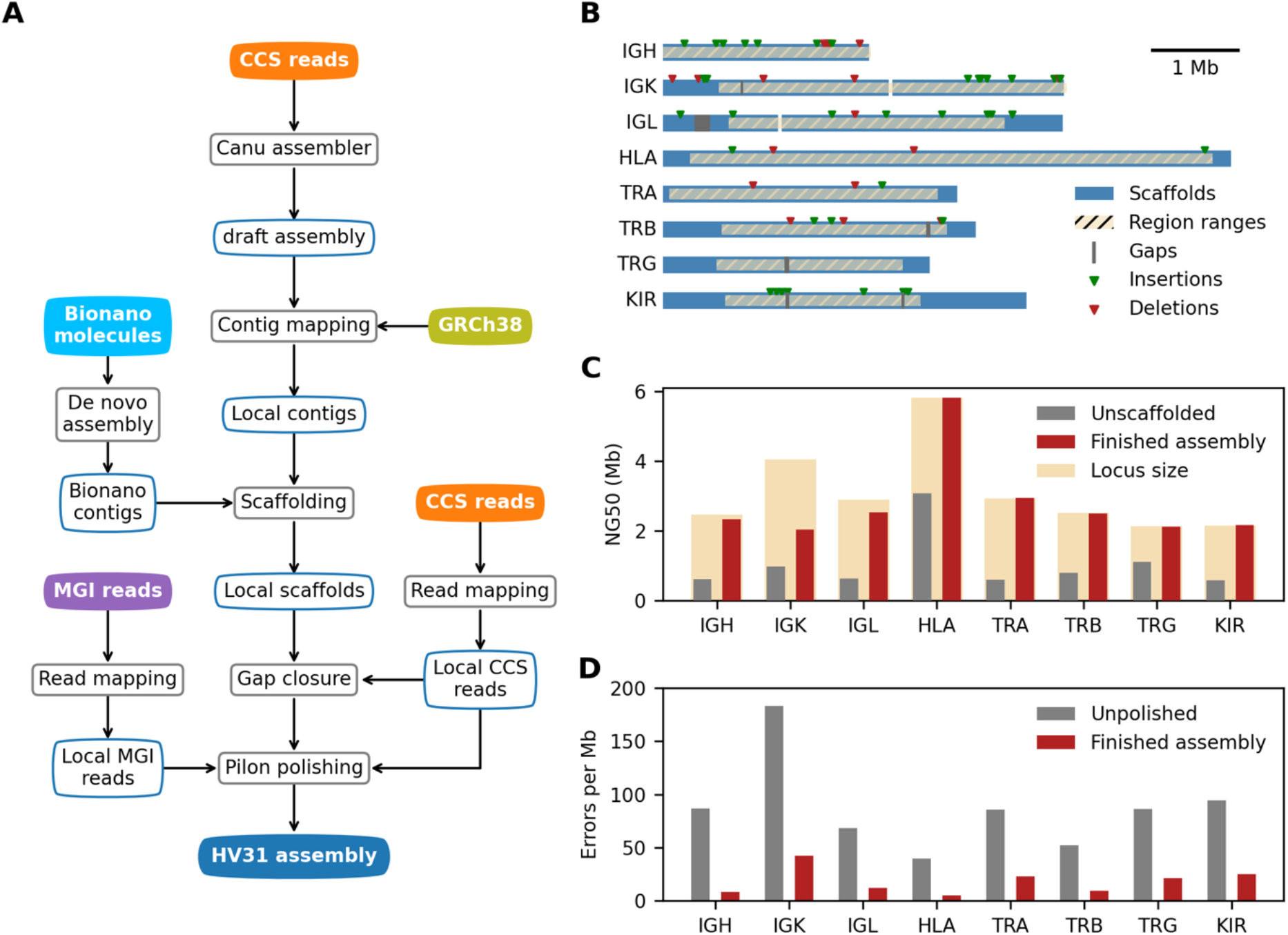
Evaluation of HV31 *de novo* assemblies. (A) Diagram of *de novo* assembly workflow. CCS, circular consensus sequencing. (B) Overview of assembled scaffolds in 8 selected regions. Heterozygous SVs on the unassembled haplotype that are larger than 1 kb in size are shown as red or green triangles. (C) Bionano scaffolding improved the continuity of the assembly, as measured by NG50, which is the length of the longest contig/scaffold that, along with longer contigs/scaffolds, covered 50% percent of each locus. (D) The polishing step reduced the number of errors in the assembly, as estimated from k-mer multiplicity in validation datasets.

#### Initial assembly

We used the Canu assembler [23] applied to CCS reads to produce an initial whole-genome assembly. We aligned the resulting contigs to GRCh38 and extracted all contigs that overlap with the predefined regions of interest, hereafter referred to as local contigs, for further processing. Local contigs were highly fragmented (Fig 1C and S1 Fig), reflecting the unusual genomic complexity in these regions. The assembly also contained multiple shorter contigs (referred to as “haplotigs” below) aligning to the same location as longer contigs in some regions, which either represent assembly errors or genuine differences between haplotypes (S1 Fig).

#### Scaffolding

We next used the local contigs with Bionano optical imaging data to produce longer continuous scaffolds. Imaged DNA molecules had an observed mean length of 149 kb, substantially longer than reads from other datasets involved in this study (S2 Fig). We used the proprietary Bionano Solve algorithm to align the local CCS contigs to contigs assembled from these imaged molecules and implemented a modified version of the BiSCoT algorithm [24] (see Methods) to order and orient the local contigs accordingly. This process also removes or merges in haplotigs that can be effectively aligned to the hybrid scaffolds. Finally, we confirmed that the remaining haplotigs represented substantial duplication of scaffolded contigs using a k-mer based method (see Methods), and removed these from downstream analysis. The scaffolds generated by this process fully covered six of the eight regions with a single scaffold, while the IGL and IGK regions were assembled with two scaffolds each (Fig 1B and 1C).

#### Gap filling and polishing

We further improved the assembly quality by carrying out a gap-closing step (which filled in nucleotide information for missing bases between adjacent contigs in a scaffold) using TGS-GapCloser [25] applied to regional CCS reads, resulting in the closing of seven gaps. We also implemented a polishing step using Pilon [26] applied to regional CCS and MGI reads, correcting erroneous bases in the assembly that likely originate from sequencing errors. To avoid bias due to read selection, for both processes we selected relevant reads using a double-alignment process that first aligns all reads to the initial whole-genome assembly, and then realigns the subset of reads mapping to local contigs to the fully scaffolded assembly (referred to as locally aligned reads below; see Methods).

#### Variant calling

We used the available long-read data to call heterozygous SVs using the HV31 assembly as reference (Fig 1B). In brief, SVs were called separately from locally aligned CCS, CLR and ONT long reads using PBSV (for CCS and CLR) and Sniffles (for CCS, CLR and ONT). A computational approach based on unique k-mers [27] was used to refine read alignment before variant calling (see Methods). Across the eight regions, 1,366 SVs were reported by PBSV or Sniffles, 491 of which were jointly supported by two or more dataset-software combinations (S3 Fig and S2 Dataset; including 179 >100 bp and 23 >1 kb in length). As a comparison point, we also created a dataset of SV calls based on 10x Genomics linked-reads sequencing data using the Long Ranger pipeline (S3 Dataset). We evaluate these calls further below.

We refer to the polished assembly scaffolds and SV dataset generated by these steps as “the HV31 assembly” hereafter; the assembly is summarized in Fig 1 and compared to the GRCh38 reference in Fig 2. As we describe below, the per-base error rate of these assembled regions is on the order of 5-50 errors per Mb (Fig 1D), which is of a similar magnitude to recently published whole-genome assemblies based on CCS data [16,28], although some structural errors do remain. We also note that six gaps remain in HV31 (denoted as dark grey bars in Fig 1B); these lie outside regions aligning to core immune system genes but could potentially be improved with additional processing.

**Fig 2.**
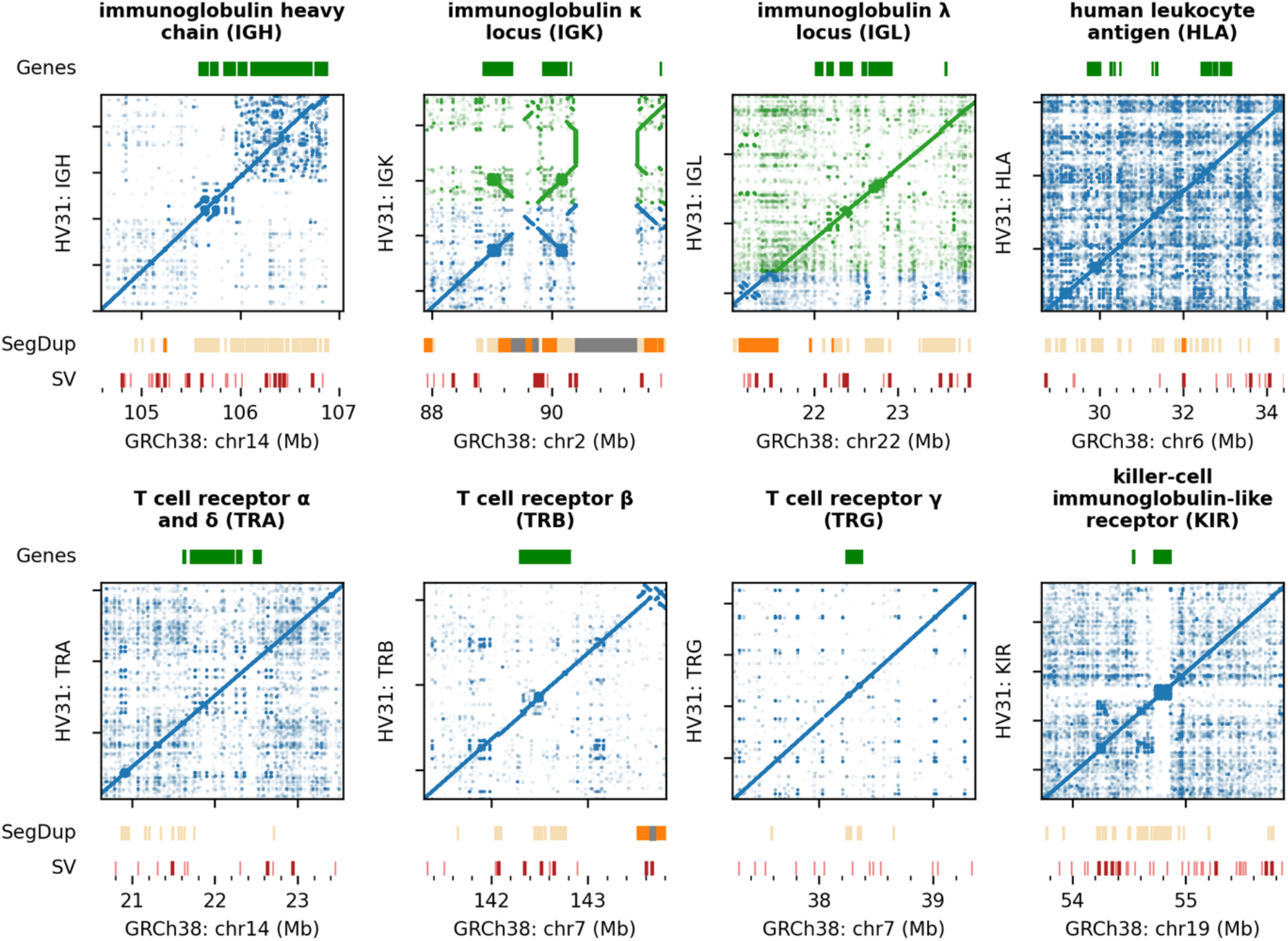
Comparing the HV31 assembly with GRCh38. Main panels show dot plots indicating locations of shared k-mers (k = 50) on the GRCh38 reference (x axis) and on the HV31 assembly (y axis) for each locus studied. In the IGK and IGL loci, the two scaffolds in the HV31 assembly are shown in blue and green, respectively. Plots are annotated as follows: Gene, core genes of each locus; SegDup, segmental duplications defined as sequence fragments that are ≥ 1 kb in length and ≥ 90% identical to another fragment (segmental duplications with identity ≥ 99% are highlighted in orange; reference gaps are shown in gray); SV, structural variants detected in the HV31 assembly relative to GRCh38. Structural variants larger than 1 kb are highlighted in dark red.

### Structural variation revealed by comparison to GRCh38

We used k-mer sharing plots (i.e. “dot plots” [29]) to compare HV31 to the GRCh38 reference sequence (Fig 2). Each point in these plots represents a k-mer (k ≤ 50) that is shared by both the reference sequence and the HV31 assembly; the observed pattern of points therefore provides a visualization of similarities and differences between the two assemblies. This comparison suggests that the HV31 assembly is relatively complete for the eight regions, without apparent missing sequence (apart from the six gaps mentioned above) or chimera sequences. HV31 contains two scaffold breaks at the IGK and IGL loci; both are located near long (≥ 100 kb) SDs that are highly identical (≥ 99%) and indicate that this type of SD remains challenging for current assembly methods. In contrast, genomic loci with higher proportions of shorter, low-similarity SDs such as the HLA and KIR were completely resolved in the HV31 assembly.

Close inspection of these plots (S4 Dataset) reveals many large (≥ 1 kb) SVs that differ between GRCh38 and the HV31 primary assembly. To systematically characterize these SVs, we aligned the assembly to GRCh38 and applied Assemblytics [30]. Assemblytics reported 145 SVs, 55 of which were ≥ 1 kb in size (Fig 1B and S1 Dataset). The majority (65.5%) of the reported SVs involved expansions or contractions of repeat elements, while the rest were insertions or deletions of unique sequences. The KIR locus harbors the highest number (29) of SVs, followed by IGH (28) and IGH (24) loci.

### Validation of assembly accuracy using unassembled sequencing reads

Given that HV31 differs structurally from GRCh38, an important question is how the structure of our assembly of these regions can be confirmed (or conversely how any remaining errors can be identified) without reliance on a reference sequence. Motivated by previous work [31,32], we adopted an approach based on computing the multiplicity of each assembly k-mer in a validation dataset, which we here take as the set of sequence reads from all low error-rate platforms in our data (S1 Table). This dataset has over 150× coverage of k-mers appearing in both copies of the genome (Fig 3A), and is sufficiently high-coverage that heterozygous k-mers, and k-mers in higher repeat numbers, can be separated from homozygous k-mers (Fig 3A and 3B). Under the assumption that sequence reads are approximately uniform across the genome, the multiplicity of each k-mer in the validation dataset should be proportional to the its copy number in the HV31 assembly [33]; any discrepancies therefore indicate heterozygous variation or assembly errors.

**Figure 3.**
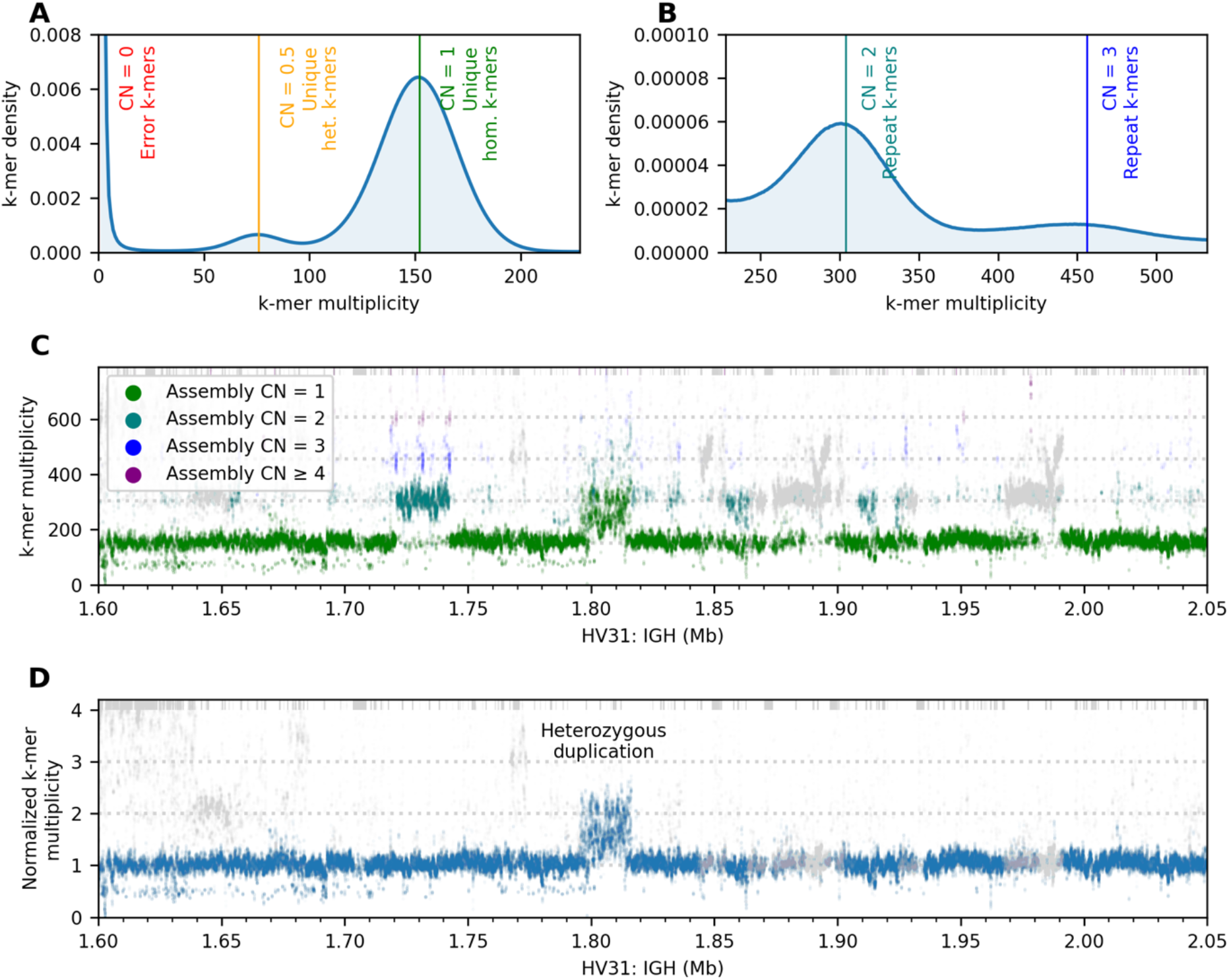
Reference-free assembly validation based on k-mer multiplicity. (A, B) Histogram of k-mer multiplicity (k = 31) in unassembled reads in the validation dataset. Vertical lines show locations of distribution peaks; text indicates interpretation of k-mers near these peaks (het., heterozygous; hom., homozygous; CN denotes assumed true copy number of the k-mer in the diploid HV31 genome). (C) k-mer multiplicity (y axis) plotted against k-mer position for a repeat-rich sequence fragment in the IGH region of the HV31 assembly. Green, blue and purple colors denote the k-mer copy number in the HV31 assembly scaffolds. Non-specific k-mers that are also found outside the IGH region are colored gray. (D) validation k-mer multiplicity normalized against assembly copy number for the same sequence fragment as in (C). In (C) and (D), k-mers with multiplicity beyond the axis limits are stacked at the top of the plots. S15 Fig and S16 Fig show panels C and D extended to all eight regions, respectively.

To leverage this, we plotted validation multiplicity in comparison to scaffold multiplicity across all regions (S15 Fig and S16 Fig; illustrated for part of the IGH region in Fig 3C and 3D). The assembly and validation dataset are in generally good agreement at most informative k-mers, including across several regions containing repeats. Base-level errors (which we identified as clusters of 22-mers with validation multiplicity < 5; see Methods) also appear to be relatively sparse; on average across regions the error rate was 18.1 per 1 Mb sequence (Fig 1D; improved from 85.3 per Mb prior to polishing). However, a number of locations show larger discrepancies between assembly and validation data (numbered regions in S16 Fig; S3 Table). We examined these in detail and found that many reflect heterozygous structural variants (illustrated in S4 Fig for a heterozygous deletion, and in Fig 3D for a complex heterozygous duplication that we discuss further below) as well as the aforementioned assembly gaps. However, a small subset of these locations indicates possible structural errors in our assembly. These include a ∼30 kb duplication in the HLA region that we confirmed is collapsed in HV31 (S4 Fig), as well as three relatively extensive stretches of elevated multiplicity in the IGK and IGL regions where we were unable to fully confirm the assembly structure using the k-mer approach. We also implemented a comparison to contigs *de novo* assembled from optical mapping data, which suggested that several of these regions were correctly assembled (see Methods). In general, this analysis indicates that our assembly of HV31 is substantially accurate apart from three repeat-rich segments in the IGK, IGL and HLA regions that may still contain errors.

We note two issues that impede validation by short k-mers. First, accurate measurement of k-mer multiplicity in highly repetitive regions is challenging; in our assembly this is particularly relevant to the IGK region which contains extensive regions of k-mers with very high copy number. A complementary approach using coverage of locally aligned ONT reads may be preferable in such regions; in our analysis this gave similar results (S16 Fig). Secondly, we noted a trend towards a drop in coverage of the validation k-mers in regions encoding T cell receptors (S15 Fig and S16 Fig). We interpret this as likely due to the use of DNA from PBMCs for some platforms included in validation (S1 Table).

### HV31 contains diverse complex immune system structural variants

As detailed above, a nontrivial number of large structural variants exist between GRCh38 and HV31 as well as between the two haplotypes of HV31. To assess the impact of these SVs on core genes, we used an alignment process to identify the best-matching variant of each immunoglobulin and T cell receptor variable gene segment, and each HLA and KIR gene within the relevant IMGT or IPD database (see Methods; Fig 4 and S5 Fig). Relative to GRCh38, the HV31 scaffolds contain both insertions and deletions of gene sequence in the IGH, IGK, IGL and TRB regions. It also contains allelic variation in all regions except TRG, and we also noted a small number of genes that differ from the best matching IMGT allele, and may represent novel sequence. We note that HV31 gene content also differs substantially from the GRCh37 assembly [21].

**Figure 4.**
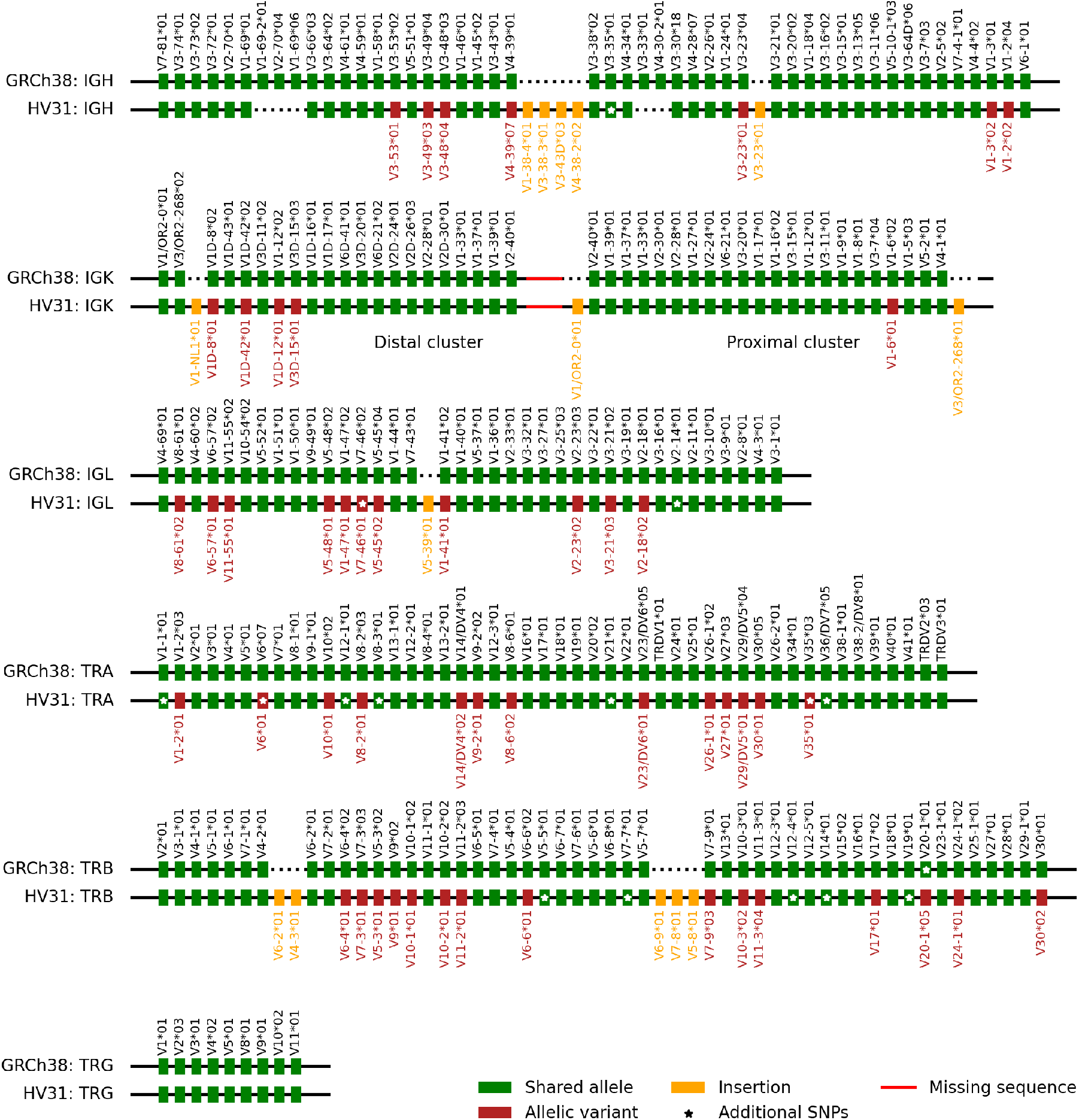
HV31 assembly content of immunoglobulin and T cell receptor variable (V) genes compared with GRCh38. Pseudogenes are not shown. V genes in each region are arranged according to their relative order on the positive-sense strand. Allelic variants refer to genes where the best-matching HV31 allele differs from the GRCh38 allele. Insertions refer to genes in the HV31 assembly that cannot be matched to a GRCh38 gene. Alleles with identical sequences, such as *TRBV6-2*01* and *TRBV6-3*01*, are not distinguished. Alleles that carry additional SNPs compared to the best-matching reference allele are marked with stars. The sequence fragment between IGK proximal and distal clusters that remains not fully resolved is denoted as a red line.

In interpreting these results, some care must be taken because of the consensus nature of the HV31 scaffolds, which do not necessarily represent a single haplotype at each locus. To elucidate underlying genetic variation, we investigated the genetic basis of the observed copy number changes in detail, focusing on the IGH and TRB regions and described in the following sections.

### A tandem repeat within a 45kb CNV involving *IGHV1-69* and *IGHV2-70*

Variation in the copy number of *IGHV1-69* and *IGHV2-70* genes has previously been reported [21]. These genes are present in three and two copies in GRCh38, respectively. In the HV31 scaffold, we found only one copy of *IGHV1-69* and *IGHV2-70* remaining, as the result of a 45 kb copy number contraction (Fig 4 and Fig 5A; variant IGH_b_29 in S1 Dataset). The earlier GRCh37 reference genome shared a similar haplotype in the IGH region, with only one copy of *IGHV1-69* and *IGHV2-70* genes. This haplotype has been suggested to be more common worldwide than the GRCh38 haplotype [14] and comparison to validation k-mers indicates it is homozygous in HV31 (S15 Fig). We noted that this CNV appears to be effectively callable by aligning reads to GRCh38, e. g. manifesting as a coverage gap in aligned PacBio CCS reads (Fig 5B); a co-located deletion was also called by the 10x pipeline, though the endpoints and length appeared inaccurate (S3 Dataset).

**Figure 5.**
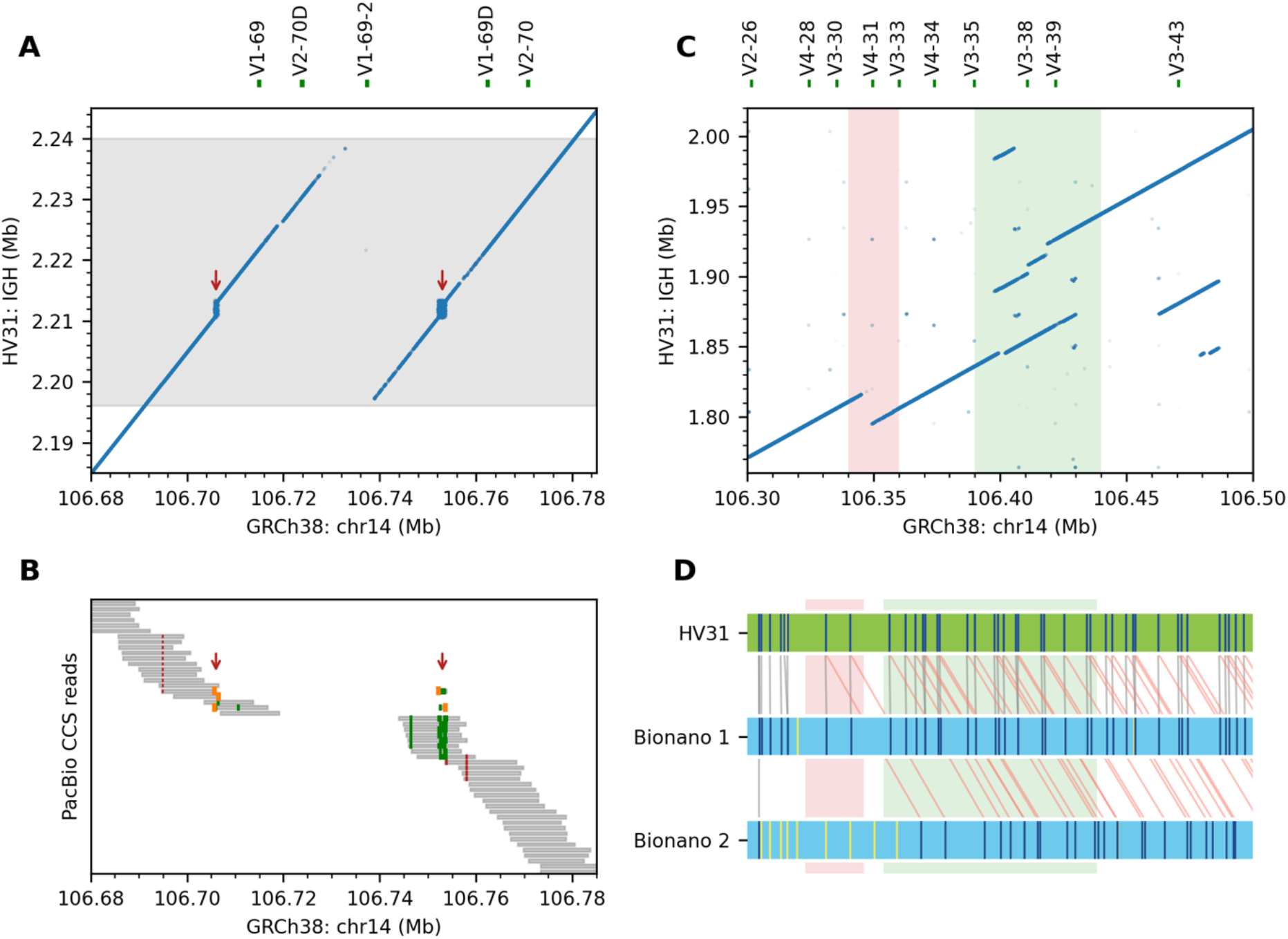
IGHV genes feature highly variable copy numbers. (A) Collapsed repeats (red arrows) in GRCh38were resolved in the HV31 assembly inside a 45 kb copy number contraction (shaded in grey). (B) Pileup of CCS reads aligned to GRCh38. The 45 kb CNV was evident from the uncovered central region. The resolved repeats were shown as alignment breakpoints (orange) or insertions (green) within the aligned segments (gray). Nearby short deletions (red) were also evident. (C) Complex structural variations found between IGHV3-21 and IGHV3-43, including a 25 kb copy number contraction (red) and an 80 kb complex duplication event (green). (D) Alignment patterns of two Bionano contigs confirm the 25 kb copy number contraction in (C), while suggesting the presence of a three-copy haplotype. Vertical lines in the HV31 assembly or Bionano contigs represent DLE-1 recognition markers. Aligned and unaligned markers are shown in dark blue and yellow, respectively.

Within this 45 kb CNV, we also noticed a 2.66 kb cluster of tandem repeats with a 59-mer motif (Fig 5A and S7 Fig) that was not correctly assembled in either GRCh37 or GRCh38 (see GenBank: AC245369.4). Similar repeat clusters have also been reported for CHM1 and NA19240 samples, though the copy numbers of the 59-mer motif varied [15].

### A compound heterozygous CNV involving *IGHV3-30*

A third prominent feature of the IGH region is the deletion of *IGHV4-30*/*IGHV4-31* and *IGHV3-33* (which is considered to be a copy of *IGHV3-30* [14]; Fig 4 and Fig 5C). Inspection of validation k-mers (Fig 3D), Bionano contigs (Fig 5D) and spanning CLR reads (S8 Fig) reveals an unusual feature: the unassembled HV31 haplotype carries an expansion relative to GRCh38 that includes three copies of these genes. As a consequence, this CNV is not associated with coverage changes in reads aligned to GRCh38. This observation is consistent with previous work [21] which reports this region as a hotspot for SVs, with the diploid copy number of *IGHV3-30* and related genes ranging from zero to six. This CNV was not called accurately by any of the SV calling methods we employed (S1-S3 Datasets).

### An 80 kb complex duplication involving multiple IGHV genes

HV31 carries additional copies of *IGHV1-38, IGHV3-43, IGHV4-38* and *IGHV3-38* genes compared to GRCh38, that are contained in a ∼80 kb duplication with complex structure (Figure 4 and Figure 5C). Inspection of k-mer multiplicity and ONT coverage depth data implies this duplication is homozygous (S15 Fig). However, this duplication was not called by any of the methods we used to call SVs (S1-S3 Datasets). We interpret this as resulting from difficulty in aligning the two sequences; consistent with this, we observed that the CCS reads in this region displayed suspicious alignment patterns when mapped to GRCh38, which were improved when mapped to the HV31 assembly (S9 Fig).

### Large insertions incorporating novel TRBV genes

In the TRB region, we detected a ∼11 kb homozygous insertion near *TRBV6-2* and another ∼19 kb insertion near *TRBV5-7* (Fig 6A). Both are supported by Bionano contigs (Fig 6B), and both insertions incorporated sequence fragments that are not found in GRCh38, with limited homology to adjacent sequences (Fig 6A). Comparison to k-mer validation again implies both insertions are homozygous. Assemblytics identified duplications at both locations but with inaccurate length and sequence content (S1 Dataset). The HV31 scaffold was consistent with an alternative contig for the TRB locus included in GRCh38 (RefSeq NG001333.2; S10 Fig). By comparing NG001333.2 with GRCh38, we confirmed that the 11 kb insertion introduced *TRBV4-3* and *TRBV6-2* genes and a *TRBV3-2* pseudogene, while the 19 kb insertion introduced the *TRBV6-9, TRBV7-8* and *TRBV5-8* genes (Fig 6 and S10 Fig).

**Figure 6.**
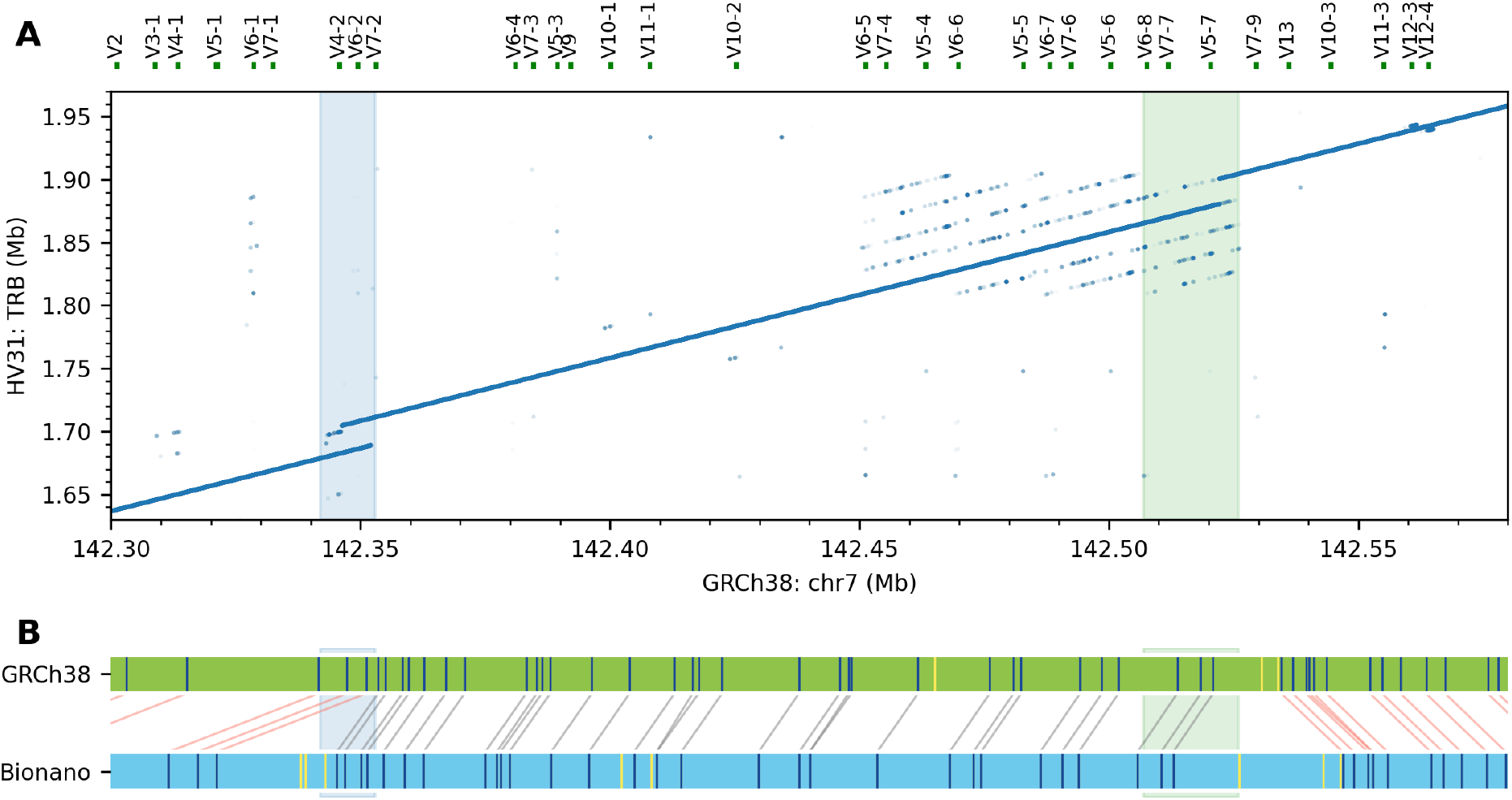
Large insertions in the TRB region. (A) HV31 harbors a 11 kb insertion (blue) and another 19 kb insertion (green) relative to GRCh38. (B) A Bionano contig (blue) aligned to GRCh38 (green), confirming the two insertions in (A).

### Reference gaps amidst complex segmental duplications resolved in the HV31 assembly

In addition to the complexities engendered by structural variation, gaps in the reference genome constitute another potential impediment to the analysis of genetic variation. Large gaps typically arise due to highly repetitive sequence that is challenging to assemble (e.g. heterochromatin regions that often consist of megabase-scale tandem satellite repeats), and their functional significance remains largely unexplored [34].

We were therefore interested to note that the HV31 assembly closes three large gaps in the GRCh38 assembly, and partly closes a fourth. Three of these gaps lie in the IGK region (S11 Fig), while the fourth lies within a 400 kb region of human-specific high-identity segmental duplications [35] ∼1Mb downstream of TRB genes. We here focus on the largest such gap, a ∼1 Mb gap annotated as heterochromatin in GRCh38, located between the distal cluster of IGK genes and the centromere of chromosome 2 (S11 Fig and S12 Fig). Examination of this gap revealed a ∼650 kb sequence assembled as an array of approximately 115 imperfect tandem copies of 6 kb repeat units (S12 Fig). Most of the repeat units contain a 22-bp signature sequence (TTCGATTCCATTTGATGATTCCAT), indicating that the heterochromatin sequence belongs to the human satellite HSat2B family [34]. The assembled heterochromatin sequence does not contain the recognition motif of DLE-1 (CTTAAG) used in generating optical maps, and we were therefore unable to directly confirm the arrangement using Bionano contigs (S11 Fig). However, assembled copy number of much of this sequence appears consistent with validation data (S16 Fig).

Notably, the HV31 assembly also contains a novel sequence fragment within the assembled heterochromatin sequence (S12 Fig). This 32 kb fragment appears unique to the region, sharing no significant homology with either the rest of the heterochromatin region nor any part of GRCh38. We found this 32 kb fragment, along with 76.8 kb flanking sequences, was over 90% identical to a 108.8 kb unplaced sequence (GenBank: AP023554.1; S12 Fig) assembled recently from individuals of Japanese ethnicity [36]. Similar ‘islands’ of unique sequence amid heterochromatin regions have previously been suggested for chromosome Y [37] and chromosome 21 [38].

Eleven heterochromatin gaps remain in GRCh38, with estimated sizes ranging from 20 kb to 30 Mb, and only the largest heterochromatin gap in chromosome X has been resolved in the form of a *de novo* assembly so far [27]. The resolved heterochromatin sequence for chromosome 2 in the HV31 assembly may provide insight into other problematic heterochromatin regions.

## Discussion

Genetic regions encoding the human immune response are among the most important for medical science, harboring determinants of both infectious and immune-mediated diseases. Despite this, the complex structure and diverse nature of many of these loci has hindered assessment of their contribution to health in large-scale genetic studies. Resolving this will likely require the creation of reference haplotype variation datasets linked to genome function. The greatest progress in this direction has been made for inference of HLA alleles [39–41] and to a lesser extent KIR alleles [42] based on large databases of known gene sequences. More recent approaches have used targeted long-read sequencing to characterize immunoglobulin variation [15]. However, much functional genetic variation affecting core immune system genes is still to be discovered and made accessible to larger studies. The decreasing cost and improving performance of long-read sequencing technologies suggests a possible route to this through the *de novo* assembly of representative individual genomes, although computational, cost, and analytic challenges remain in practice.

In this study, we contribute to this program by assembling eight of the most complex immune system regions in a single healthy European individual identified as HV31, who was recruited as part of a larger study of genomic and transcriptomic variation in immune cell types. To do this, we based our assembly on DNA extracted from CD14^+^ monocytes, allowing us to assess the germline haplotype configurations in the immunoglobulin and T cell receptor regions. We exploited accurate PacBio CCS data for the initial assembly, and used optical mapping data to scaffold the assembled contigs, followed by additional gap closing and polishing steps (Fig 1). Alongside this, we also developed a set of methods to validate assembly correctness, enabling us to establish with reasonable confidence the accuracy of our assembly. Although some structural errors do remain, and may be resolvable with further work, our results suggest we have produced essentially correct representations of seven of eight regions studied here, with some caveats in the IGK region. Our assembly thus adds to growing numbers of reference-quality assemblies that can be utilized in these regions [15,22,43] and to the catalogue of known alleles at these loci.

An ideal description of a diploid sample would involve two fully assembled contigs for every somatic chromosome, each representing one of the two inherited haplotypes. This approach is sought by several emerging experimental and computational approaches [44–46]. Generating a phased assembly inevitably involves a trade-off among cost, phasing accuracy and assembly continuity, which is further complicated by the presence of large high-identity duplications [46]. Given the limited depth of CCS reads in our data (S1 Table), we instead chose to represent the HV31 genome as a set of consensus (i.e. mixed-haplotype [47]) scaffolds supplemented with a list of heterozygous variants (S2 Dataset). This is a pragmatic approach which also simplifies downstream analysis as the assembly can be directly used in place of a reference genome without additional preparation.

Fully assembling regions containing long, complex repeat structures from shotgun sequencing remains a challenging problem; any approach must somehow distinguish reads coming from different repeat copies but identify those that truly overlap – while allowing for sequencing errors. Exemplifying this challenge, our assembly of the IGK region fills a ∼1 Mb heterochromatin gap in GRCh38 that largely consists of high copy-number repeats but also contains unique sequence (S12 Fig). Direct confirmation of the assembled structure of IGK is difficult using our data and this may require further experimental methods as comparison to validation data (S15 and S16 Fig) is inconclusive due to the high repeat number, but our assembly of this region is compatible with other recently reported assemblies. The fact that most of the repetitive regions within the loci studied here are correctly assembled is of interest in itself, as it implies that the repeats are old enough (or divergent enough) to be effectively distinguished.

Given the high diversity observed in these regions it is unsurprising that HV31 carries structural variants which make it differ from current genome builds and from other previously reported samples. Some of the variation we have identified in HV31 has previously been associated with phenotypes - for example, copy number polymorphism of *IGHV1-69* is known to strongly correlate with the prevalence of this gene in expressed antibody repertoires, which is preferentially used in antibodies against certain influenza strains and the HIV-1 virus [7]. However, we have also highlighted extensive variant haplotypes that have not previously been reported. The degree of variation observed in this single sample indicates that much haplotype variation of immunoglobulins and T cell receptor regions remains to be discovered. It is also notable that many of the more complex variants we have identified are not accurately called by the variant calling methods we employed (S1-S3 Datasets). Thus, both studies with larger sample sizes and further methodological improvements will likely be needed to fully elucidate variation in these important regions.

## Methods

### Ethics statement

HV31 was recruited as a healthy volunteer under approval by the Oxfordshire Research Ethics Committee (COREC reference 06/Q1605/55).

### DNA extraction, sequencing and optical mapping

Blood sample of a healthy female donor of European ancestry identified as HV31 was used in this study. For PacBio CCS and CLR sequencing, Oxford Nanopore sequencing, 10x linked-read sequencing and MGI stLFR linked-read sequencing, the DNA was extracted from CD14^+^ monocytes isolated from PBMC using CD14 antibody-conjugated beads. DNA extraction was performed with QIAGEN MagAttract HMW DNA kit following manufacturer’s instructions, with slight adaptations. In brief, 220 µl Buffer ATL and 20 µl Proteinase K were added to a suspension of 1×10^6^ CD14^+^ monocytes. The mixture was incubated overnight at 56 **°C**, shaking at 900 rpm to lyse the cells. The lysate was then processed according to manufacturer’s instructions for the purification of high-molecular-weight genomic DNA from tissue.

For MGI standard short-read, MGI coolMPS and Illumina PCR-free sequencing, the DNA was extracted from PBMC with NEB Monarch Genomic DNA Purification Kit (T3010) following manufacturer’s instructions. Library preparation, sequencing and optical mapping were performed following the instructions of the respective platform providers.

### Region definition

Eight genomic regions encoding key components of the human immune system, including HLA, IG, TCR and KIR were selected for investigation (S1 Table). Each region was defined as a core range in GRCh38 that contained genes related to immune system components, with additional flanking sequences added to both sides. The core range were typically selected based on the respective reference sequences in the NCBI RefSeq database [18]. As exceptions, the core range of the HLA region was defined as the genomic range from *GABBR1* to *KIFC1* genes [19], and the KIR region was defined as the genomic range from *KIR3DL3* to *KIR3DL2* genes [20]. The flanking sequence was typically 1 Mb on either side. As exceptions, the telomeric flanking sequence in the IGH region was limited to 164 kb by the length of chromosome 14. In addition, we expanded the centromeric flanking sequence in the IGK region by 0.67 Mb to bridge a 1 Mb heterochromatin gap present in GRCh38.

### Whole-genome *de novo* assembly

Canu v1.9 [23] was used to perform whole-genome *de novo* assembly for HV31 based on PacBio CCS reads, with the following parameters: -pacbio-hifi <CCS_FASTQ> genomeSize=3235000000 -minInputCoverage=1 -stopOnLowCoverage=1. The resulting contigs were mapped to GRCh38 using minimap2 [48] with the following parameters: -ax asm5 --secondary=no. Contigs that mapped to the 8 loci of interest were extracted as local contigs.

### Hybrid scaffolding and haplotig removal

Hybrid scaffolding was performed using Bionano Solve, a proprietary software provided by Bionano Genomics (https://bionanogenomics.com/), with default parameters. We used a custom script based on BiSCoT [24] to improve the contiguity and quality of the resulting scaffolds. Specifically, we merged adjacent contigs in a scaffold if they overlap with each other, as inferred from shared enzymatic labelling sites or sequence alignment. If the two adjacent contigs were expected to be non-overlapping, then the size of the gap between them was estimated based on the distance of nearest labelling sites. In addition, we incorporated shorter contigs into longer ones if the shorter contig represented a subsequence of the longer contig, and aligned better with the Bionano genome maps.

After scaffolding, we removed duplicated contigs or scaffolds that presumably represent alternative haplotypes (‘haplotigs’) using a custom k-mer based method. In brief, we listed all unique 22-mers for each contig or scaffold and compare these sets of 22-mers in a pairwise manner. If a shorter contig had more than 80% of unique 22-mers shared with a longer contig, then the former was considered as a haplotig and removed from the assembly.

### Read mapping

Sequencing reads from each locus of interest were required for various purposes including gap closing, polishing, error rate estimation and assembly validation based on alignment coverage and patterns. In order minimize reference bias, we first mapped the reads from each sequencing dataset using minimap2, and then extracted reads that mapped to contigs that represent each locus of interest [49]. The extracted reads were again mapped with minimap2 to the scaffolded or finalized assembly as appropriate for specific applications.

A unique k-mer anchoring method [27] was used to improve the mapping of long reads in repetitive regions. In brief, given a set of locus-specific reads and a corresponding reference sequence, we first defined a set of anchoring k-mers for each locus of interest. Only k-mers that appeared to be unique in both short read sequencing datasets (31 ≤ multiplicity ≤ 231) and the reference sequence (copy number = 1; no occurrence outside the locus) were selected as anchoring k-mers. Then, we mapped the reads to the reference with minimap2 using parameters -n 50 -r 10000, which enabled the output of up to 50 alignments for each read, with gap sizes up to 10 kb in each alignment. An optimal alignment for each read were was then selected based on the number of bases shared with the reference that were part of an anchoring k-mer. These selected alignments were pooled into a new BAM file, after filtering out alignments that were shorter than 5 kb. The resulting BAM file were used for polishing and reference-free alignment validation.

### Gap closing and polishing

Gap closing was performed using TGS-GapCloser v1.0.1 [25] with PacBio CCS reads. Sequencing reads were first mapped to the whole genome assembly produced by Canu, which enabled locus-specific read extraction. The extracted reads were used as input for TGS-GapCloser, which was executed using the following parameters: -ne --tgstype pb --g_check. Polishing was performed using Pilon [26] with CCS reads and MGI paired-end short reads extracted in a similar manner. The default parameters were used. For clarity, the finalized scaffolds were displayed and coordinated based on the relative order and orientations of the corresponding sequence in GRCh38 in visualization steps.

### Error rate estimation and reference-free assembly validation

Jellyfish [50] was used to count the multiplicity of each k-mer (k = 22 or 31) from a pooled FASTQ dataset of PacBio CCS, MGI standard short-read, MGI CoolMPS, MGI stLFR linked read, 10x Linked-Read and Illumina PCR-free sequencing platforms (S1 Table), with the following parameters: jellyfish count -m <k> -s 30G --min-qual-char “?” -C. The accumulated sequencing depth of the pooled FASTQ dataset was 262×. In each read, k-mers that include bases with base quality < 20 were excluded. For error rate estimation, k-mers (k = 22) in the HV31 assembly with multiplicity < 5 were classified as erroneous k-mers, and clustered by their positions in the assembly, allowing a maximum of k - 1 correct k-mers between two adjacent erroneous k-mers in each cluster. The number of erroneous k-mer clusters per Mb assembled sequence was used as an indicator of the error rate of the HV31 assembly.

For reference-free assembly validation, we define the normalized multiplicity (N) of each k-mer (k = 31) in the HV31 assembly as N = M / (C × D), where M is the multiplicity of that k-mer in the validation dataset, C is the copy number of that k-mer in the HV31 assembly, and D is the mode multiplicity of unique homozygous k-mers in the validation dataset, as estimated from the k-mer multiplicity histogram (Fig 3A). The normalized k-mer coverage was visualized against the position of the k-mer, along with the normalized coverage of ONT reads aligned to the assembly using the k-mer anchoring method. Regions where the normalized k-mer coverage or normalized ONT coverage deviated from 1 were labelled and inspected for potential assembly errors (S3 Table).

### Variant calling

PBSV (https://github.com/PacificBiosciences/pbsv), a subprogram of SMRT tools was used to call heterozygous SVs from CCS and CLR reads with default parameters. Sniffles [51] was used to call heterozygous SVs from CCS and CLR and ONT reads with the following parameters: -s 3 -q 20 --ccs_reads --min_het_af 0.2 (CCS), -s 8 -q 20 --min_het_af 0.2 (CLR), or -s 15 -q 20 --min_het_af 0.2 (ONT). Unique k-mer anchoring was applied prior to SV calling. SVmerge, a subprogram of SVanalyzer [52] was used to cluster and merge SV records from output VCF files of PBSV and Sniffles, with default parameters.

### Gene variant detection

Reference variant sequences of IGHV, IGKV, IGLV, TRAV, TRDV, TRBG and TRGV genes were downloaded from the IMGT reference directory [53] (http://www.imgt.org/download/V-QUEST/IMGTV-QUESTreference_directory.zip). Reference variant sequences of HLA genes were downloaded from the IPD-IMGT/HLA database [54] (ftp://ftp.ebi.ac.uk/pub/databases/ipd/imgt/hla/fasta/hla_gen.fasta). Reference variant sequences of KIR genes were downloaded from the IPD-KIR database [55] (ftp://ftp.ebi.ac.uk/pub/databases/ipd/kir/fasta/KIR_gen.fasta). The reference gene variant sequences were mapped to GRCh38 or the HV31 assembly using Minimap2 with the following parameters: -a -w1 -f1e-9. We extracted subsequences in regions where at least one reference gene was mapped, with 20 bp flanking sequence at either side. These query sequence fragments were submitted to NCBI IgBLAST [56] (for IGHV, IGKV, IGLV, TRAV, TRDV, TRBV and TRGV genes) or NCBI BLAST+ [57] (for HLA and KIR genes) to search for matching sequences in the relevant databases, with default parameters. The top hit variant with the highest match score returned by NCBI IgBLAST or NCBI BLAST+ were assigned to each query fragment. Query fragments shorter than the top hit variant were considered to represent partial alignment and discarded.

## Data Availability

A complete list of data generated in this study (S1 Table) will be available through the EGA.

## Author contributions

### Conceptualization

Antony J. Cutler, Gavin Band, Julian C. Knight, John A. Todd. Rachael Bashford-Rogers.

### Data curation

Jia-Yuang Zhang, Hannah Roberts, David S. C. Flores, Justin P. Whalley, Gavin Band.

### Investigation

Jia-Yuan Zhang, Hannah Roberts, David Flores, Antony J. Cutler, Andrew C. Brown, Olga Mielczarek, Justin P. Whalley, Barbara Xella, Karen Oliver, Craig Corton, Emma Betteridge, Gavin Band.

### Methodology

Antony J. Cutler, Andrew C. Brown, Jia-Yuan Zhang, Gavin Band.

### Formal analysis

Jia-Yuan Zhang, Hannah Roberts, David S. C. Flores.

### Supervision

Helen Lockstone, David Buck, Antony J. Cutler, Rachael Bashford-Rogers, Gavin Band, Julian C. Knight, John A. Todd

### Visualization

Jia-Yuan Zhang.

### Writing – original draft preparation

Jia-Yuan Zhang, Gavin Band

### Writing – review & editing

Jia-Yuan Zhang, Gavin Band, Antony J. Cutler, Andrew C. Brown, Justin P. Whalley, Rachael Bashford-Rogers, Julian C. Knight, John A. Todd.

### Funding acquisition

Julian C. Knight, John A. Todd.

## Acknowledgements

The work of JYZ and JAT was supported by the Juvenile Diabetes Research Fund [5-SRA-2015-130-A-N], [4-SRA-2017-473-A-N]; the Wellcome [107212/Z/15/Z]; [203141/Z/16/Z]. Computation used the Oxford Biomedical Research Computing (BMRC) facility, a joint development between the Wellcome Centre for Human Genetics and the Big Data Institute supported by Health Data Research UK and the NIHR Oxford Biomedical Research Centre. Financial support was provided by the Wellcome Trust Core Award Grant Number 203141/Z/16/Z. The views expressed are those of the author(s) and not necessarily those of the NHS, the NIHR or the Department of Health. JYZ was supported by the China Scholarship Council-University of Oxford Scholarship. GB is a member of the MalariaGEN resource centre, supported by Wellcome [204911/Z/16/Z]. The funders had no role in study design, data collection and analysis, decision to publish, or preparation of the manuscript.

## Supporting Information

**S1 Fig. Whole-genome assembly contigs aligned to GRCh38 in each of the 8 selected loci**.

**S2 Fig. Read/molecule length distribution of PacBio CCS, PacBio CLR, ONT and Bionano optical mapping datasets involved in this study**. Red and grey vertical lines denote the N50 and mean read/molecule length for each dataset, respectively.

**S3 Fig. Supplementing the HV31 assembly with alignment-based SV calling**. A given SV call is classified as an insertion if the alternative haplotype is longer than the reference haplotype, or a deletion if otherwise.

**S4 Fig. Detecting heterozygous SVs and assembly errors from k-mer multiplicity and coverage depth patterns**. (A) A 63.9 kb heterozygous deletion in the HLA locus is revealed by reduced ONT coverage depth (orange) and k-mer multiplicity (blue). (B) A collapsed duplication in the HLA locus is revealed by elevated ONT coverage depth (orange) and k-mer multiplicity (blue). Non-specific k-mers that are also found outside the IGH region are shown in gray. In (A) and (B), k-mers with multiplicity beyond the axis limits are stacked at the top of the plots.

**S5 Fig. Haplotypes of HLA and KIR genes of HV31 compared with GRCh38. G**enes in each region are arranged according to their relative order on the positive-sense strand. Allelic variants refer to genes where the HV31 allele differ from the GRCh38 allele in sequences. Alleles that carry additional SNPs on the basis of the corresponding reference allele are marked with stars.

**S6 Fig. Schematic examples of sequence duplications and structural variations demonstrated with k-mer sharing plots**. X, Y and Z denote sequence fragments that are different from each other. Y’ denotes the reverse complement of Y. In each panel, the reference sequence is depicted on the x axis and the alternate sequence is depicted on the Y axis. The size of each structural variation can be estimated from the distance between relevant breakpoints on the plot.

**S7 Fig. GRCh38 and GRCh37 represent different IGH haplotypes**. (A) Similar to HV31, GRCh37 has only one copy of IGHV1-69 and IGHV2-70 genes. The unresolved repeats are highlighted with red arrows. Gray shade marks the position of the 45 kb CNV in HV31 relative to GRCh38 (see Figure 3A). (B) Schematic representation of GRCh38, GRCh37 and HV31 near *IGHV1-69* and *IGHV2-70* genes. Fragment R denotes the unresolved duplication which was assembled in HV31.

**S8 Fig. A compound heterozygous CNV involving *IGHV3-30***. (A) k-mer sharing plot comparing GRCh38 with itself, highlighting the two copies (blue and green) of a 21.2 kb duplication unit that harbored *IGHV3-30* and *IGHV3-33*, respectively. k = 50. (B) k-mer sharing plot comparing CLR read 101124000/2432040434 with GRCh38, demonstrating a one-copy haplotype. k = 20 (C) k-mer sharing plot comparing CLR read 92801871/034335 with GRCh38, demonstrating a three-copy haplotype. k = 20.

**S9 Fig. Misalignment resulting from large structural rearrangements in the IGH locus**. (A) k-mer sharing plot comparing the HV31 assembly with GRCh38, highlighting the 80 kb insertion between *IGHV3-37* and *IGHV7-40* which introduced extra copies of several gene fragments. k = 50. The region inspected in (B) and (C) was highlighted in blue. (B) Misalignment was observed when aligning PacBio CCS reads to GRCh38. (C) PacBio CCS reads were correctly assembled when using the HV31 assembly as reference.

**S10 Fig. The HV31 assembly is consistent with NCBI RefSeq NG_001333.2**. TRBV genes not included in GRCh38 are highlighted in green.

**S11 Fig. Three gaps flanked by high-identity repeats were filled in the HV31 assembly**. (A) k-mer sharing plot comparing GRCh38 with the HV31 assembly. k = 50. The 2.56 Mb scaffold and the 1.97 Mb scaffold in the HV31 assembly are shown in blue and green, respectively. Coverage of ONT reads aligned to GRCh38 is displayed above, and the proximal and distal clusters are annotated. Gaps in GRCh38 are shaded in gray. Novel sequence junctions in the HV31 assembly are annotated with red arrows. Sequence fragments of which extra copies were introduced in the HV31 assembly to fill in the gaps between IGK proximal and distal gene clusters in GRCh38 are highlighted in yellow; corresponding read coverage peaks confirm increased genome multiplicity of these fragments. (B) Alignment of Bionano contigs (blue) to the 2.56 Mb scaffold in the HV31 assembly (green). The approximate sequence region that maps to the GRCh38 gaps between IGK proximal and distal gene clusters is shaded in gray. For clarity, corresponding positions in the HV31 assembly in (A) and (B) were labelled with red arrows. (C) Alignment of BioNano contigs (blue) to the 1.97 Mb scaffold in the HV31 assembly (green). Approximate sequence region that maps to the GRCh38 heterochromatin gap is shaded in gray.

**S12 Fig. The heterochromatin gap in the IGK locus was filled with 650 kb complex repeat sequence**. (A) k-mer sharing plot comparing the HV31 assembly with itself in the IGK heterochromatin region. k = 50. Purple lines show the occurrence of a 22 bp HSat2B repeat signature sequence (TTCGATTCCATTTGATGATTCCAT). A 32 kb unique sequence fragment is highlighted in blue. (B) Dot plot in (A) zoomed in to reveal details of the unique sequence fragment and repeat structure. (C) Same as Figure 4C, zoomed in to show that the 32 kb unique fragment (blue) contains a DLE-1 recognition label that was confirmed by Bionano contigs. (D) k-mer sharing plot comparing the HV31 assembly (y axis) with GenBank AP023554.1 (x axis). k = 50.

**S13 Fig. ONT coverage depth in the IGK heterochromatin region**. Compared with the Minimap2 mapping (yellow), the read mapping method based on unique k-mer anchoring (green) yielded more uniform coverage depth in the highly repetitive IGK heterochromatin region.

**S14 Fig. The HV31 assembly is consistent with chr7KZ208912v1fix but harbors an inversion. (A)** k-mer sharing plot comparing chr7*KZ208912v1*fix with GRCh38, highlighting the genomic position corresponding to the 140 kb inversion in the HV31 assembly (green) and the 50 kb gap in GRCh38 (gray). k = 50. **(B)** The HV31 assembly is consistent with chr7*KZ208912v1*fix except for the 21.9 kb gap (brown) and the 140 kb inversion (green).

**S15 Fig. Assembly validation based on k-mer multiplicity**. k = 31.

**S16 Fig. Assembly validation based on normalized k-mer multiplicity and normalized ONT coverage depth**. k = 31.

**S1 Table. Summary of sequencing and genome mapping datasets**.

**S2 Table. Bioinformatics tools used in this study**.

**S3 Table. Potentially problematic regions in the HV31 assembly as identified from k-mer multiplicity**

**S1 Dataset. SVs in the HV31 assembly as detected by Assemblytics**.

**S2 Dataset. Heterozygous SVs reported by SVmerge**.

**S3 Dataset. SVs detected in the 10x Genomics Long Ranger pipeline**.

**S4 Dataset. k-mer sharing plots comparing the HV31 assembly with GRCh38 in each of the eight regions of interest**. k = 50.

