## Supplementary material for "Using *de novo* assembly to identify structural variation of complex immune system gene regions": S1 Table.docx

| Name | Cell type | Coverage | Read length |
| --- | --- | --- | --- |
| PacBio Circular Consensus Sequencing (CCS) | CD14^+^ monocytes | 12.3× | Mean = 12.7 kb; N50 = 12.7 kb |
| PacBio Continuous Long Reads (CLR) | CD14^+^ monocytes | 35× | Mean = 15.4 kb; N50 = 25.9 kb |
| Oxford Nanopore PromethION (ONT) | CD14^+^ monocytes | 63× | Mean = 8.7 kb; N50 = 10.9 kb |
| Bionano DLS optical mapping | PBMC | 152.7× | Mean = 149.4 kb; N50 = 216.4 kb |
| MGI standard short-read sequencing | PBMC | 56.8× | 100 bp paired-end |
| MGI coolMPS sequencing | PBMC | 56.9× | 100 bp paired-end |
| MGI stLFR linked-read sequencing | CD14^+^ monocytes | 51.3× | 100 bp paired-end |
| 10X Linked-Read sequencing | CD14^+^ monocytes | 40.2× | 150 bp paired-end |
| Illumina PCR-free sequencing | PBMC | 44.2× | 151 bp paired-end |
