## Supplementary material for "Using *de novo* assembly to identify structural variation of complex immune system gene regions": S2 Table.docx

| Name | Version | Reference |
| --- | --- | --- |
| Assemblytics | 1.2.1 | Nattestad, M., & Schatz, M. C. (2016). Assemblytics: A web analytics tool for the detection of variants from an assembly. Bioinformatics, 32(19), 3021–3023. https://doi.org/10.1093/bioinformatics/btw369 |
| Bionano Solve | 3.5.1_01142020 | https://bionanogenomics.com/support/software-downloads/ |
| Canu | 1.9 | Koren, S., Walenz, B. P., Berlin, K., Miller, J. R., Bergman, N. H., & Phillippy, A. M. (2017). Canu: Scalable and accurate long-read assembly via adaptive k -mer weighting and repeat separation. Genome Research, 27(5), 722–736. https://doi.org/10.1101/gr.215087.116 |
| Jellyfish | 2.3.0 | Marçais, G., & Kingsford, C. (2011). A fast, lock-free approach for efficient parallel counting of occurrences of k-mers. Bioinformatics, 27(6), 764-770. |
| Minimap2 | 2.14 | Li, H. (2018). Minimap2: pairwise alignment for nucleotide sequences. Bioinformatics, 34(18), 3094-3100. |
| MUMmer | 4.0.0beta2 | Delcher, A. L., Phillippy, A., Carlton, J., & Salzberg, S. L. (2002). Fast algorithms for large-scale genome alignment and comparison. Nucleic acids research, 30(11), 2478-2483. |
| Pilon | 1.23 | Walker, B. J., Abeel, T., Shea, T., Priest, M., Abouelliel, A., Sakthikumar, S., Cuomo, C. A., Zeng, Q., Wortman, J., Young, S. K., & Earl, A. M. (2014). Pilon: An Integrated Tool for Comprehensive Microbial Variant Detection and Genome Assembly Improvement. PLoS ONE, 9(11), e112963. https://doi.org/10.1371/journal.pone.0112963 |
| Samtools | 1.1 | Li, H., Handsaker, B., Wysoker, A., Fennell, T., Ruan, J., Homer, N., ... & Durbin, R. (2009). The sequence alignment/map format and SAMtools. Bioinformatics, 25(16), 2078-2079. |
| SMRT Tools (PBSV) | 7.0.1.66975 | https://github.com/PacificBiosciences/pbsv |
| Sniffles | 1.0.12a | Sedlazeck, F. J., Rescheneder, P., Smolka, M., Fang, H., Nattestad, M., von Haeseler, A., & Schatz, M. C. (2018). Accurate detection of complex structural variations using single-molecule sequencing. Nature Methods, 15(6), 461–468. https://doi.org/10.1038/s41592-018-0001-7 |
| SVanalyzer (SVmerge) | 0.36 | Zook, J. M., Hansen, N. F., Olson, N. D., Chapman, L. M., Mullikin, J. C., Xiao, C., Sherry, S., Koren, S., Phillippy, A. M., Boutros, P. C., Sahraeian, S. M. E., Huang, V., Rouette, A., Alexander, N., Mason, C. E., Hajirasouliha, I., Ricketts, C., Lee, J., Tearle, R., … the Genome in a Bottle Consortium. (2019). A robust benchmark for germline structural variant detection [Preprint]. Genomics. https://doi.org/10.1101/664623 |
| TGS-GapCloser | 1.1.1 | Xu, M., Guo, L., Gu, S., Wang, O., Zhang, R., Fan, G., Xu, X., Deng, L., & Liu, X. (2019). TGS-GapCloser: Fast and accurately passing through the Bermuda in large genome using error-prone third-generation long reads [Preprint]. Bioinformatics. https://doi.org/10.1101/831248 |
