## Supplementary material for "Using *de novo* assembly to identify structural variation of complex immune system gene regions": S3 Table.docx

| Region | Number | Evidence | Speculation |
| --- | --- | --- | --- |
| IGH | 1 | CLR reads | Misalignment. |
| IGH | 2 | CCS reads | Heterozygous deletion. |
| IGH | 3 | CLR reads | Heterozygous duplication; see S8 Fig. |
| IGK | 4 | HV31 assembly | Assembly gap. |
| IGK | 5 | Bionano contigs and CCS reads | Misalignment. |
| IGK | 6 | - | Unclear. |
| IGK | 7 | - | Unclear. |
| IGK | 8 | - | Unclear. |
| IGK | 9 | ONT reads | Heterochromatin microsatellite array; see S12 Fig |
| IGK | 10 | - | Unclear. |
| IGL | 11 | Bionano contigs | Misalignment; possible assembly error outside the IGL region. |
| IGL | 12 | Bionano contigs | Heterozygous deletion. |
| HLA | 13 | CCS reads | Heterozygous deletion; see S4 Fig. |
| HLA | 14 | CCS reads | Assembly error (collapsed duplications); see S4 Fig. |
| TRB | 15 | Bionano contigs and CCS reads | Heterozygous duplication |
| TRB | 16 | CLR reads | Misalignment. |
| TRB | 17 | HV31 assembly | Assembly gap. |
| TRB | 18 | Bionano contigs and HV31 assembly | Misalignment due to assembly gap (see number 17). |
| TRG | 19 | HV31 assembly | Assembly gap. |
| KIR | 20 | HV31 assembly | Assembly gap. |
