## Supplementary figures and images for "Using *de novo* assembly to identify structural variation of complex immune system gene regions"

### S1 Fig.png

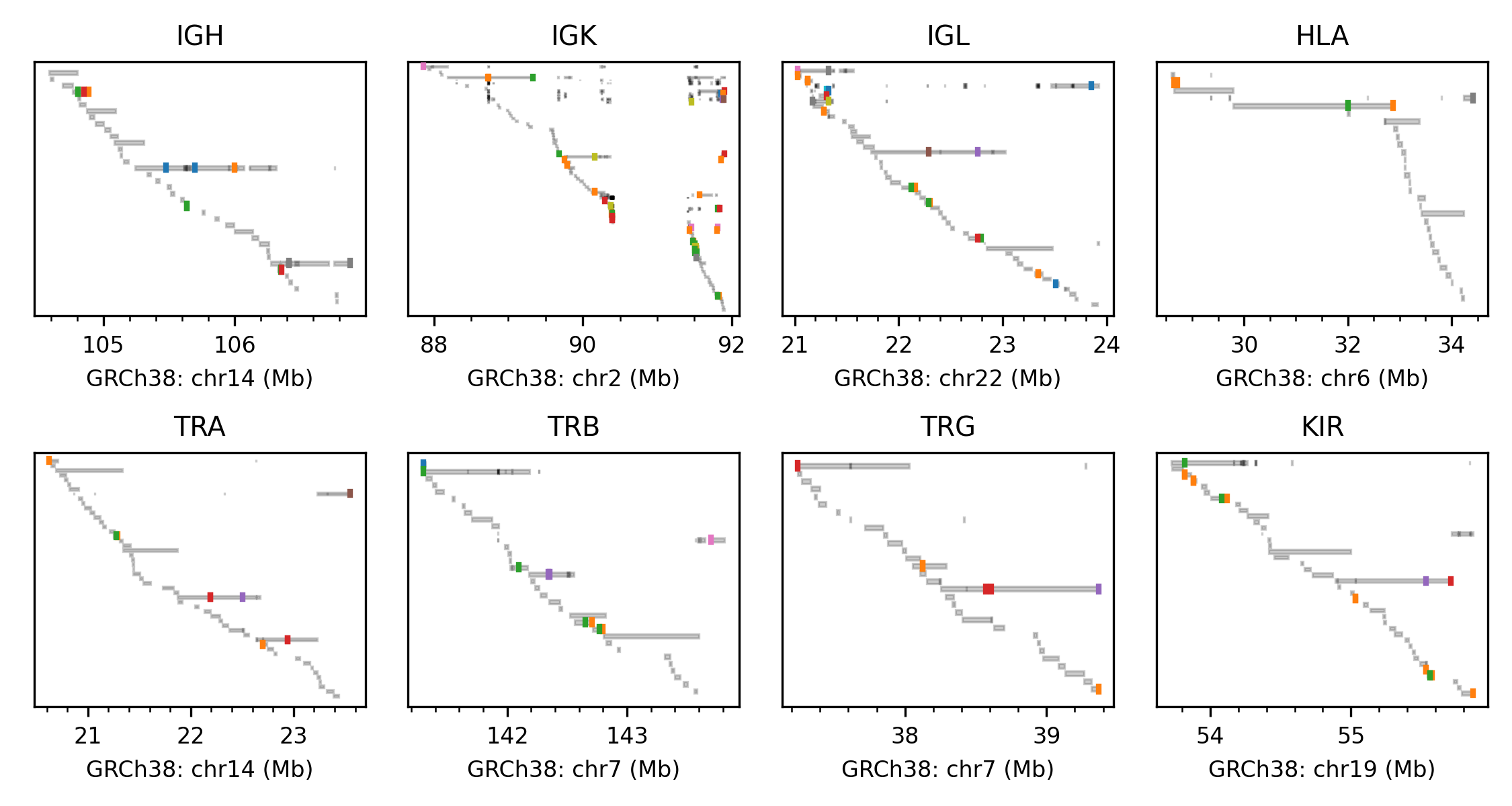

### S2 Fig.png

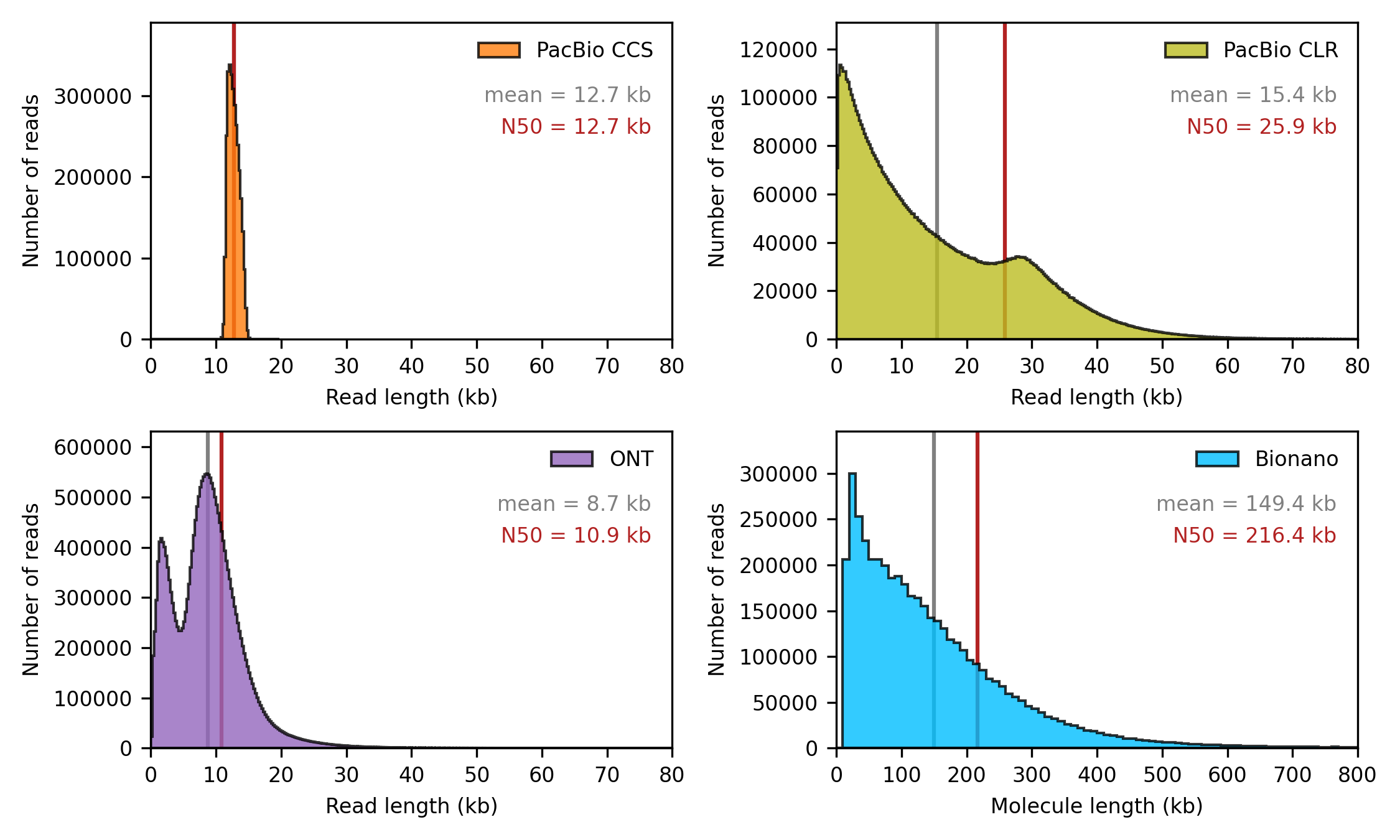

### S3 Fig.png

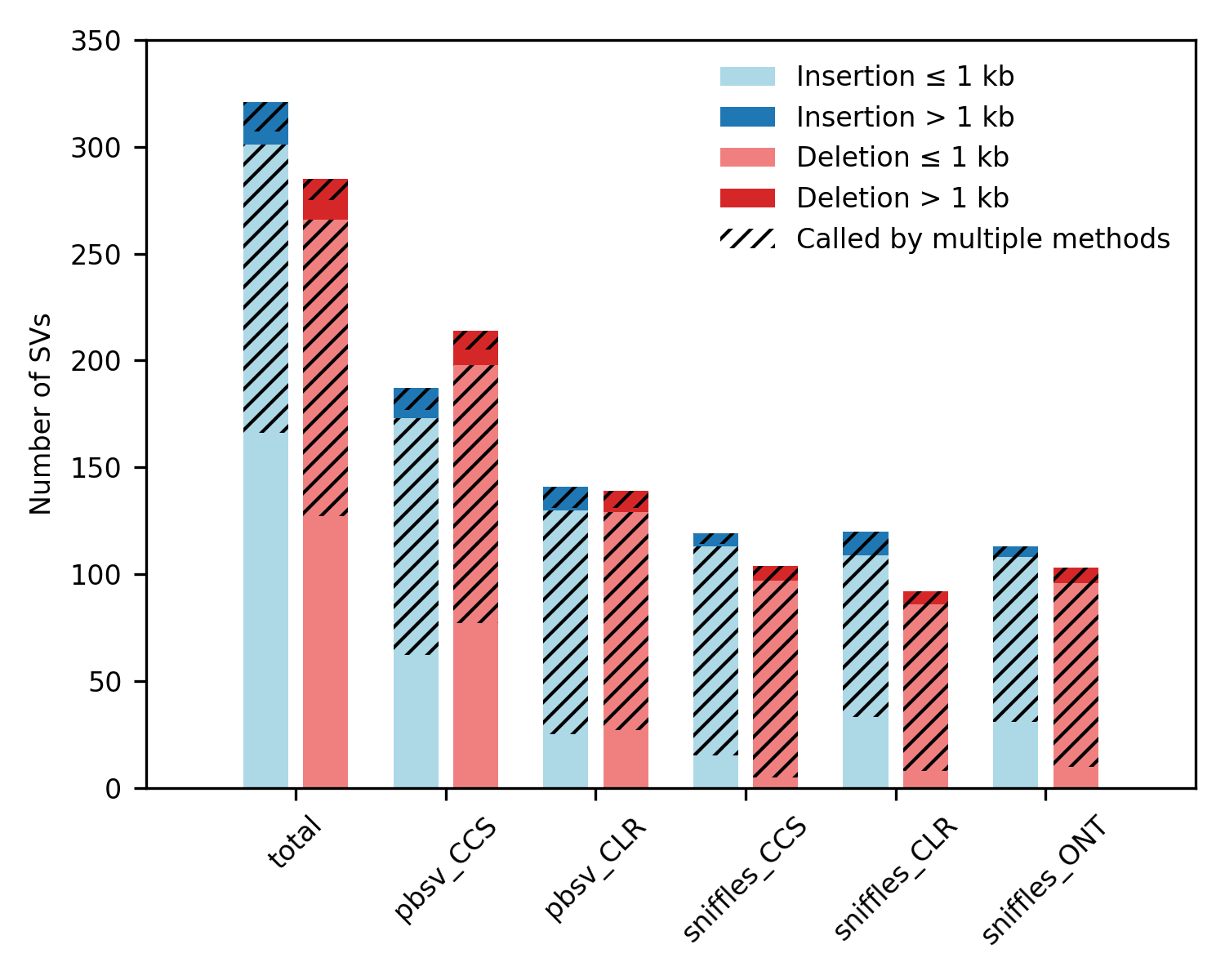

### S4 Fig.png

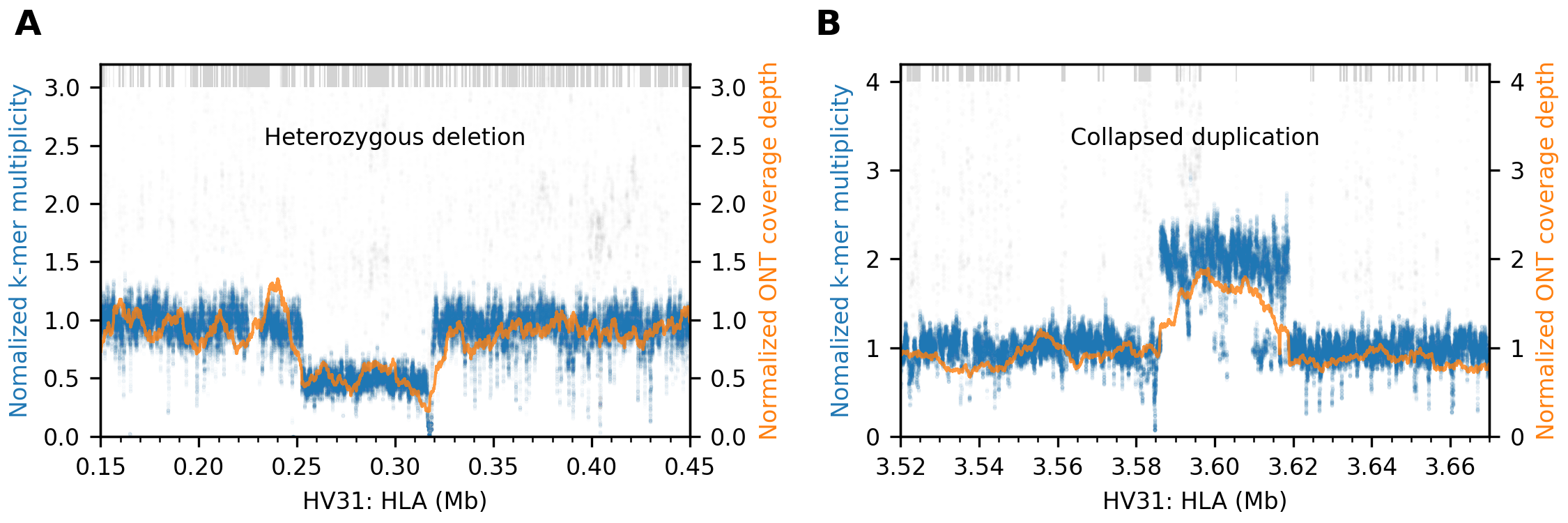

### S5 Fig.png

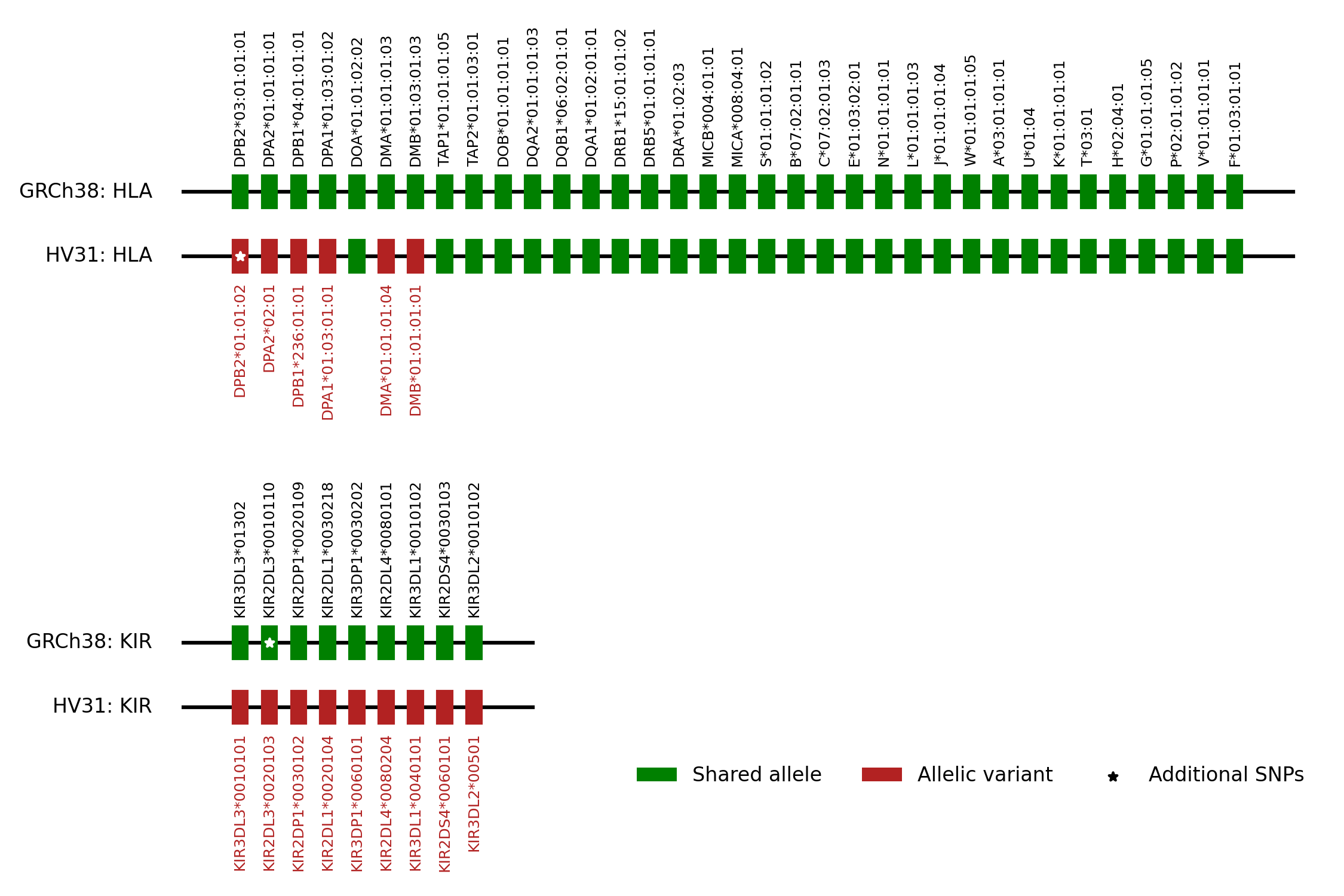

### S6 Fig.png

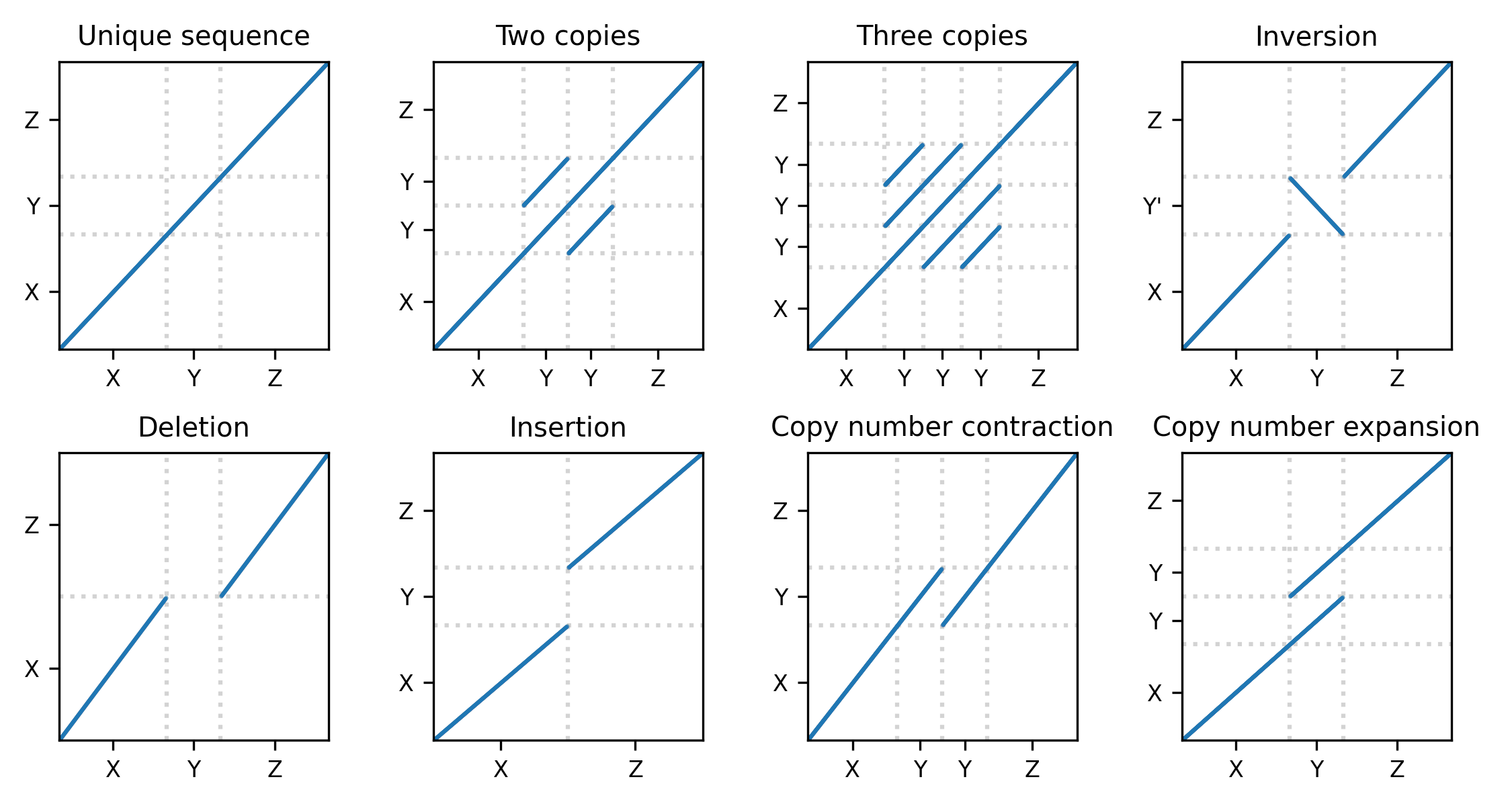

### S7 Fig.png

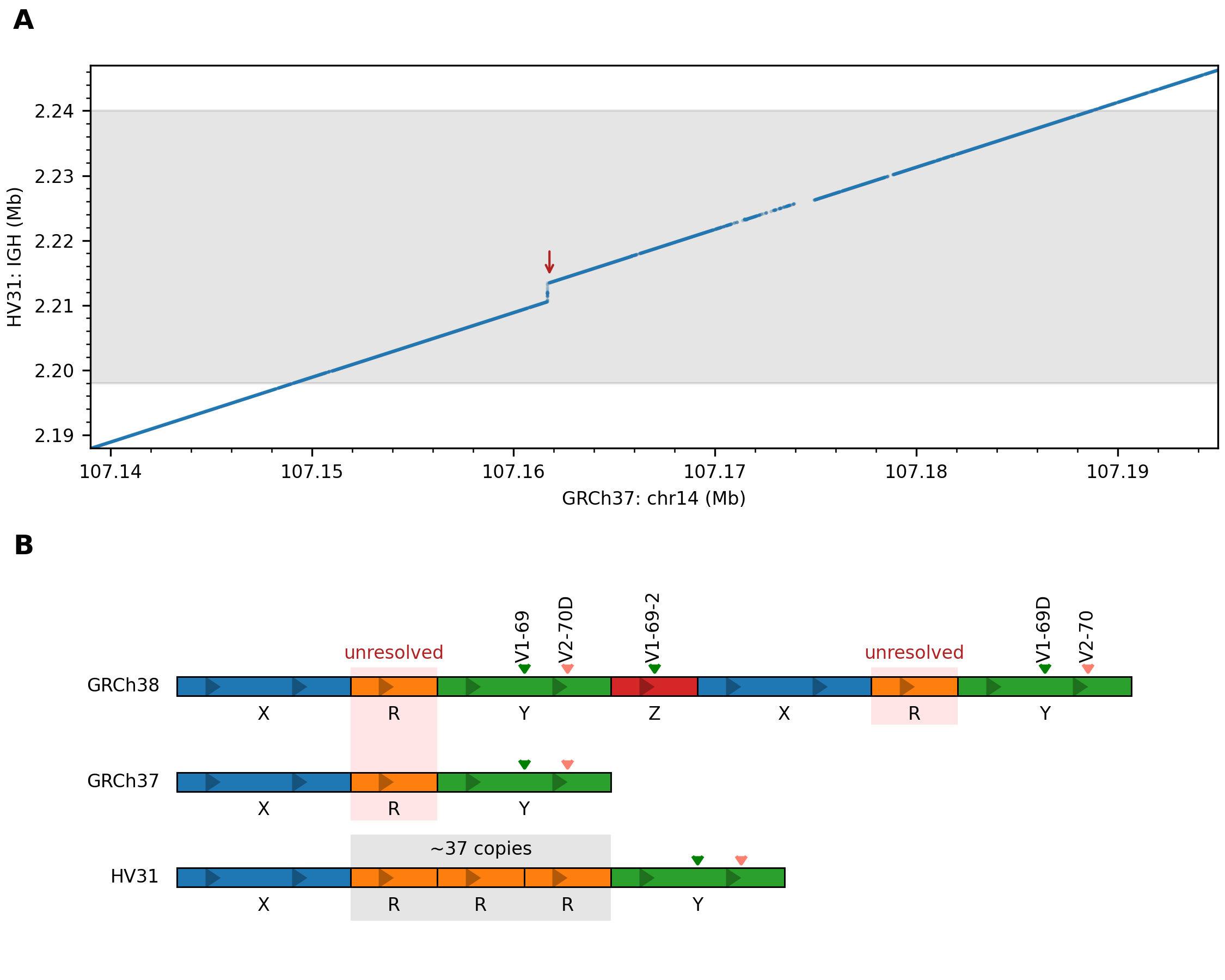

### S8 Fig.png

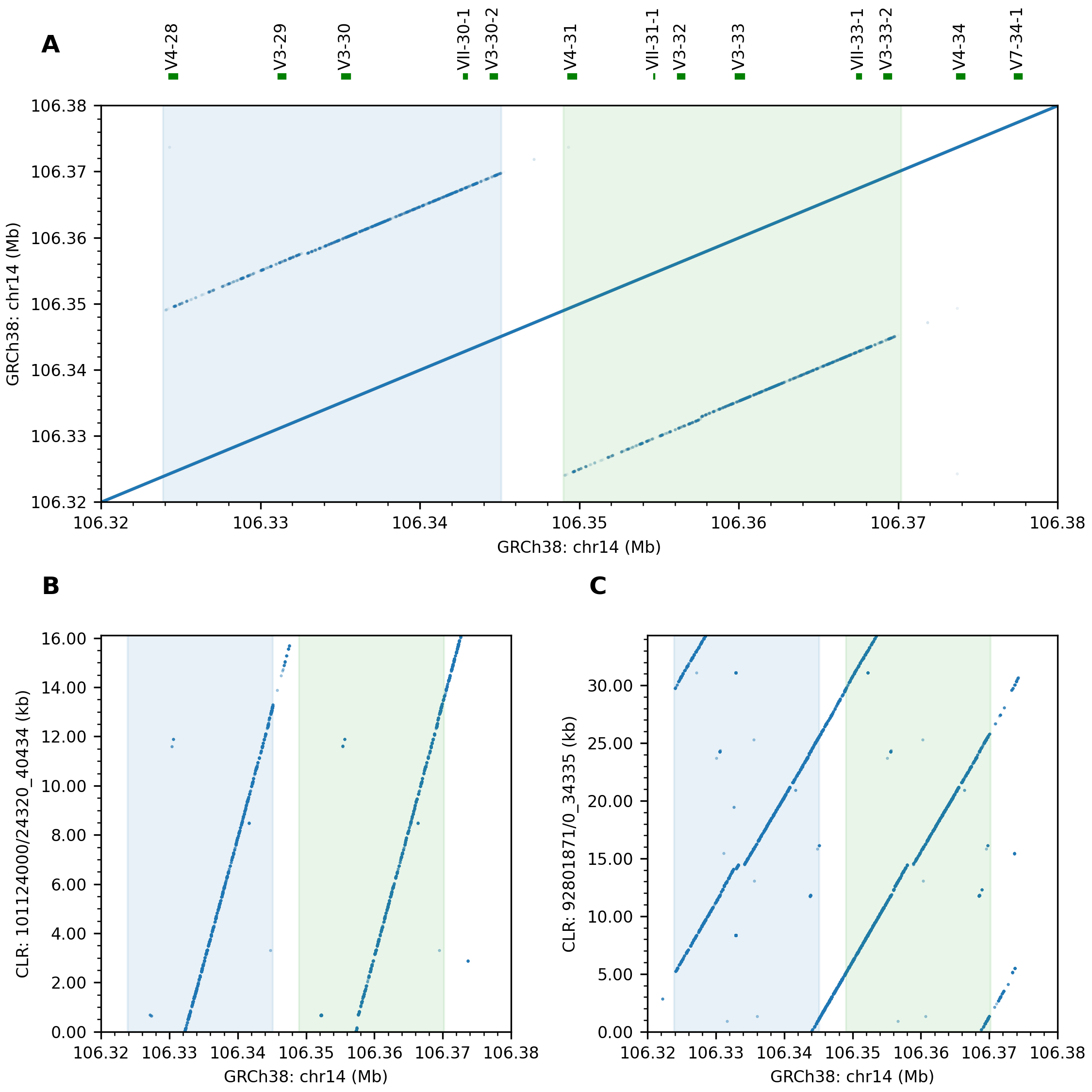

### S9 Fig.png

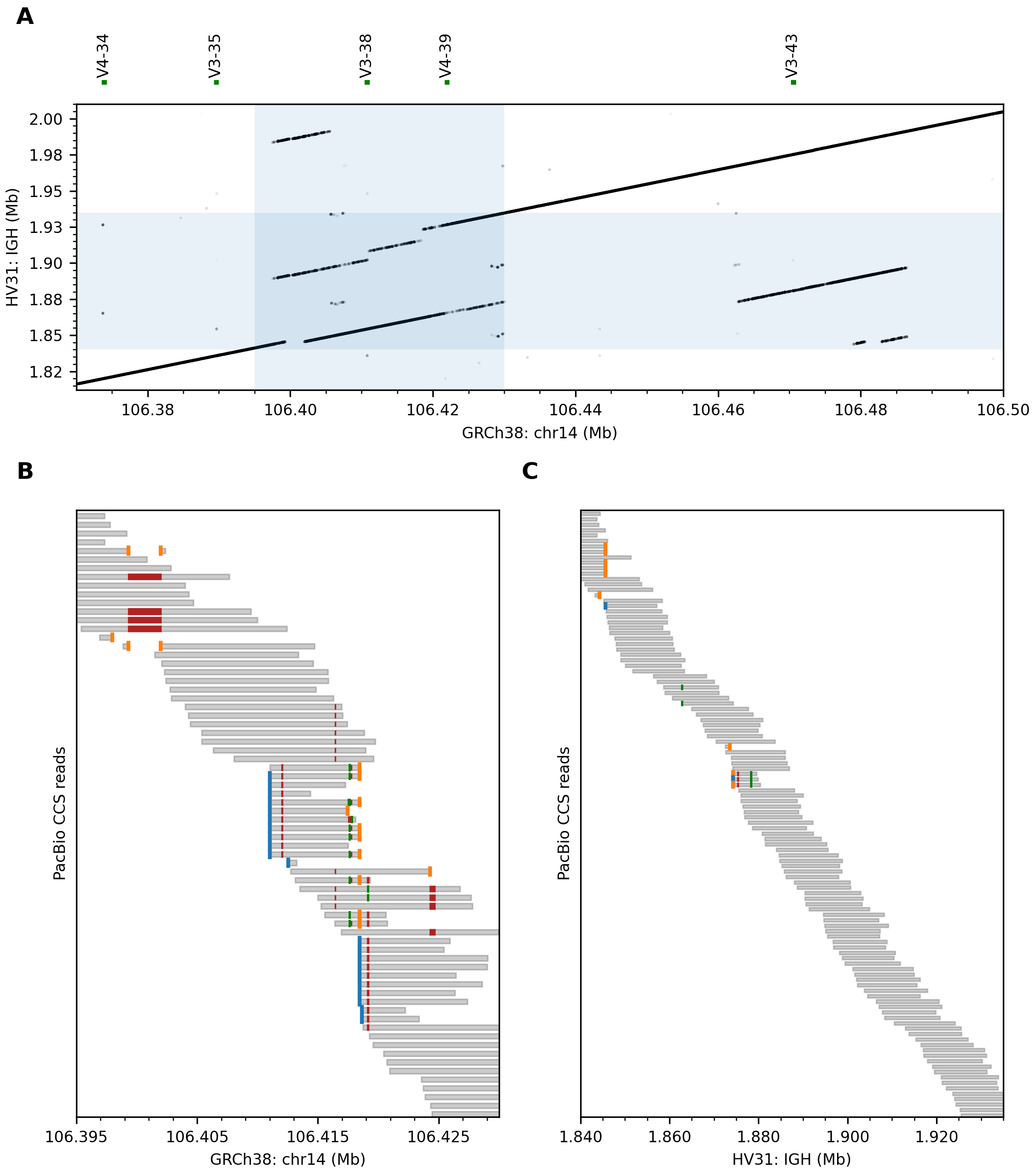

### S10 Fig.png

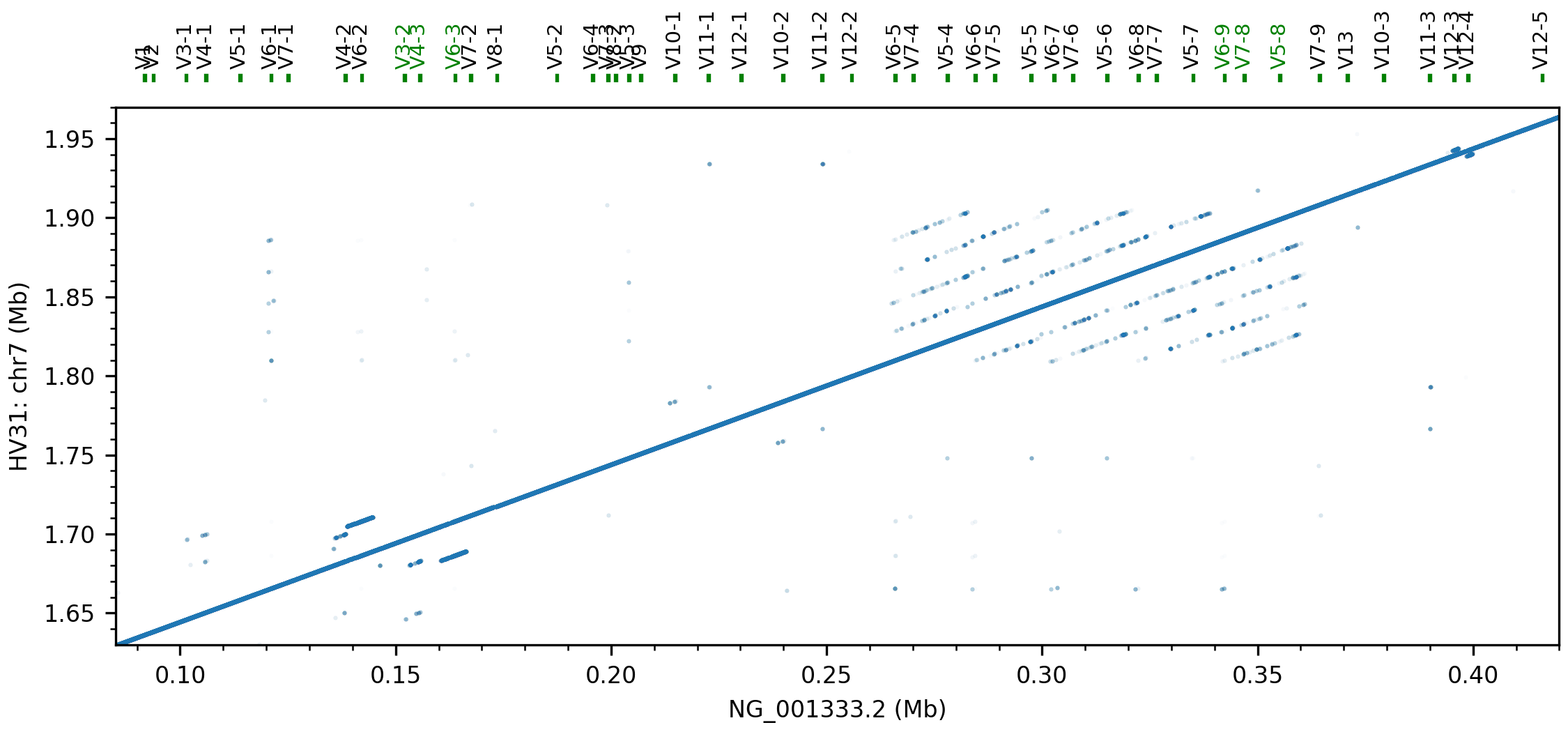

### S11 Fig.png

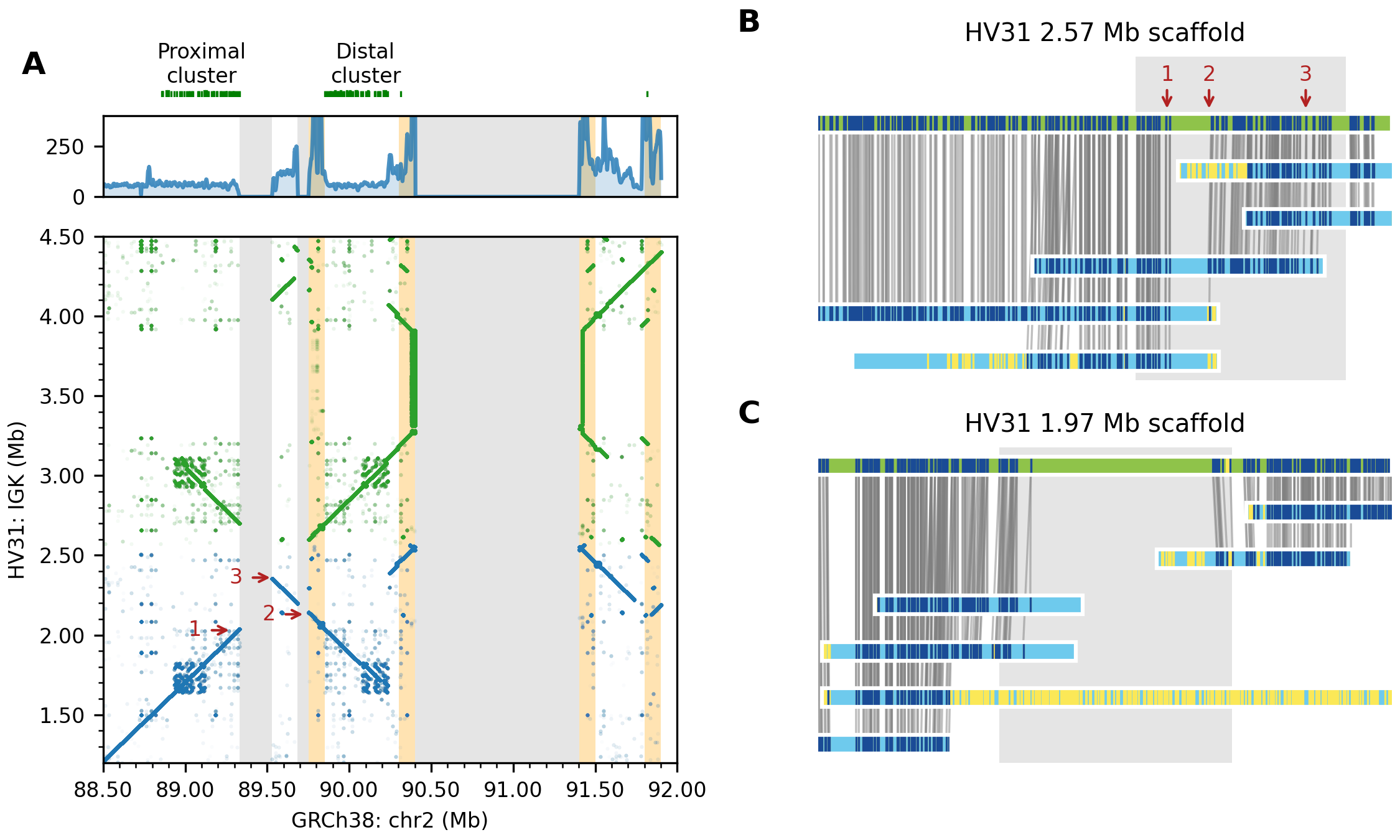

### S12 Fig.png

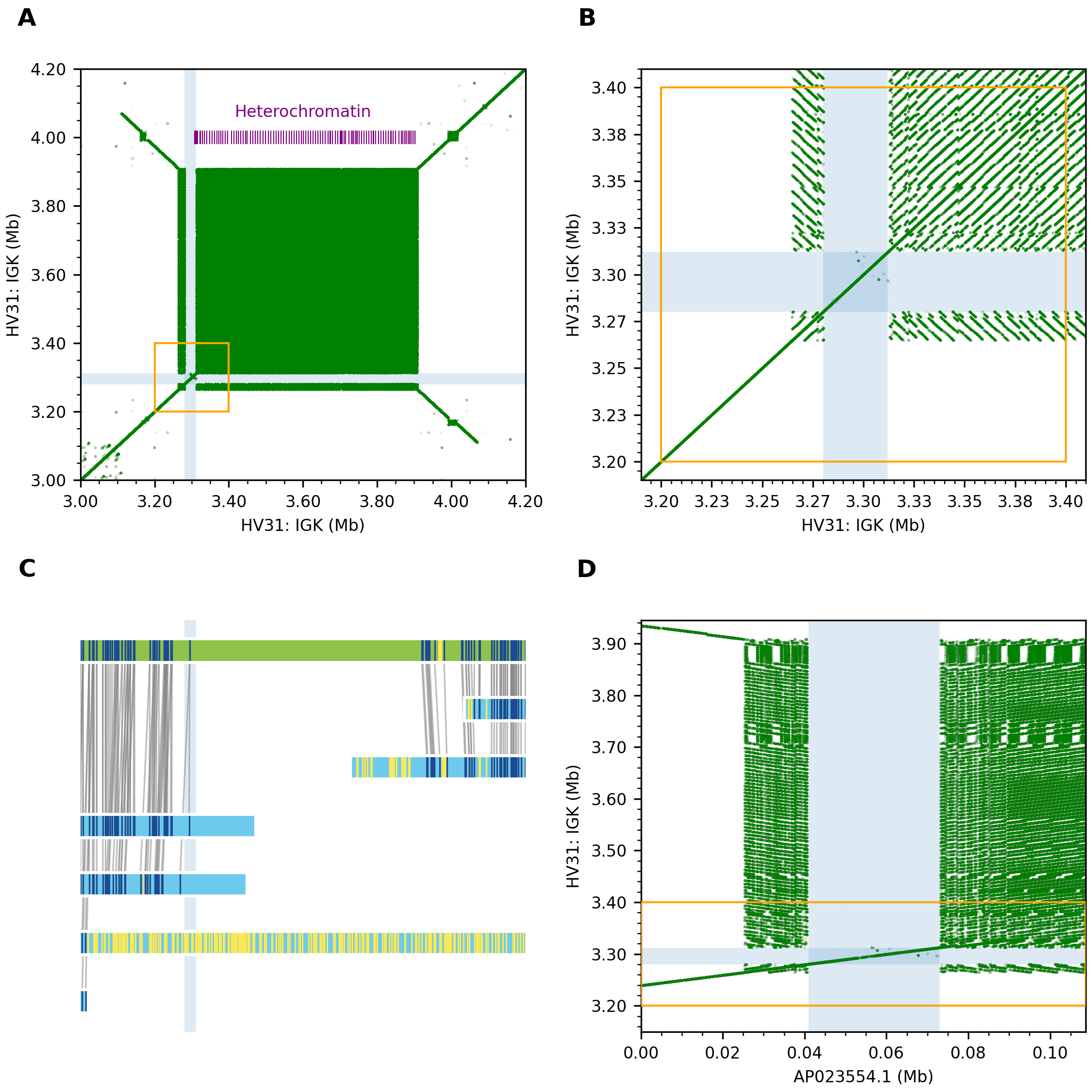

### S13 Fig.png

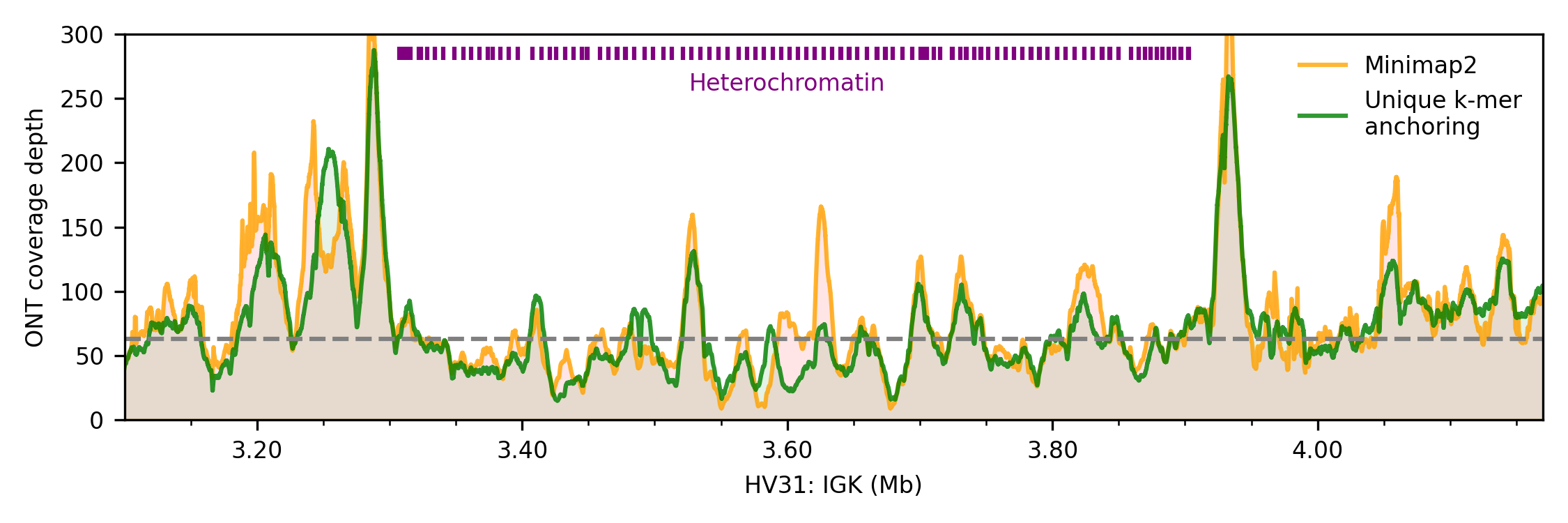

### S14 Fig.png

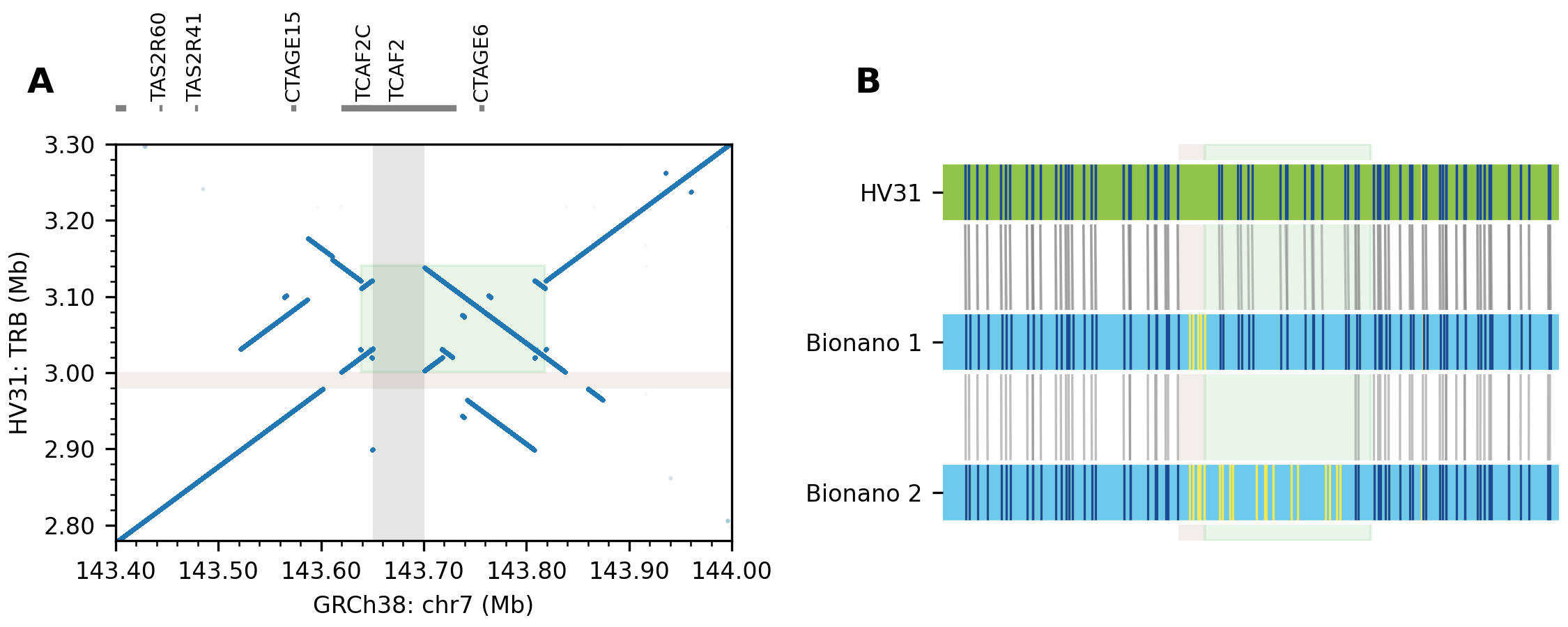

### S15 Fig.png

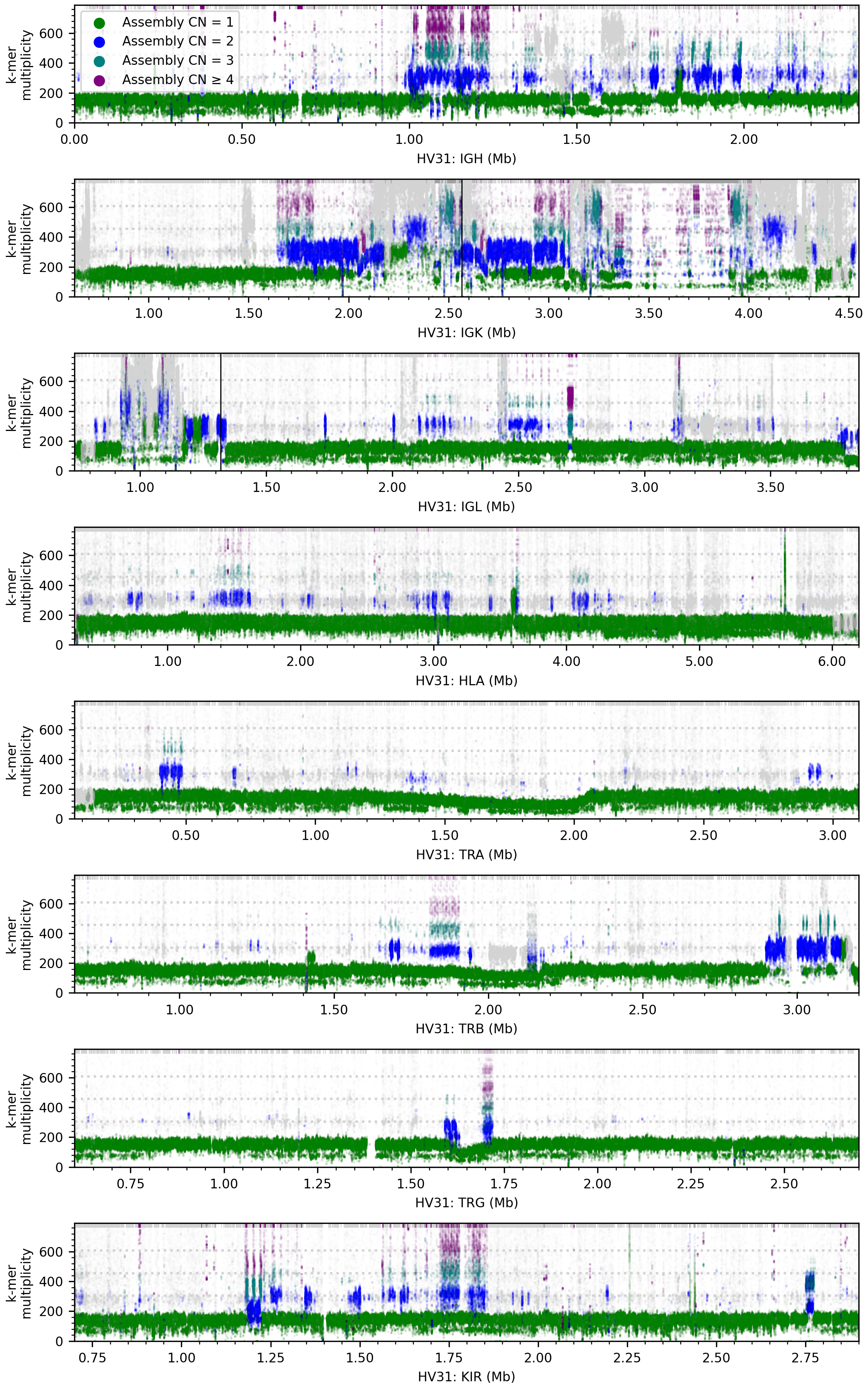

### S16 Fig.png

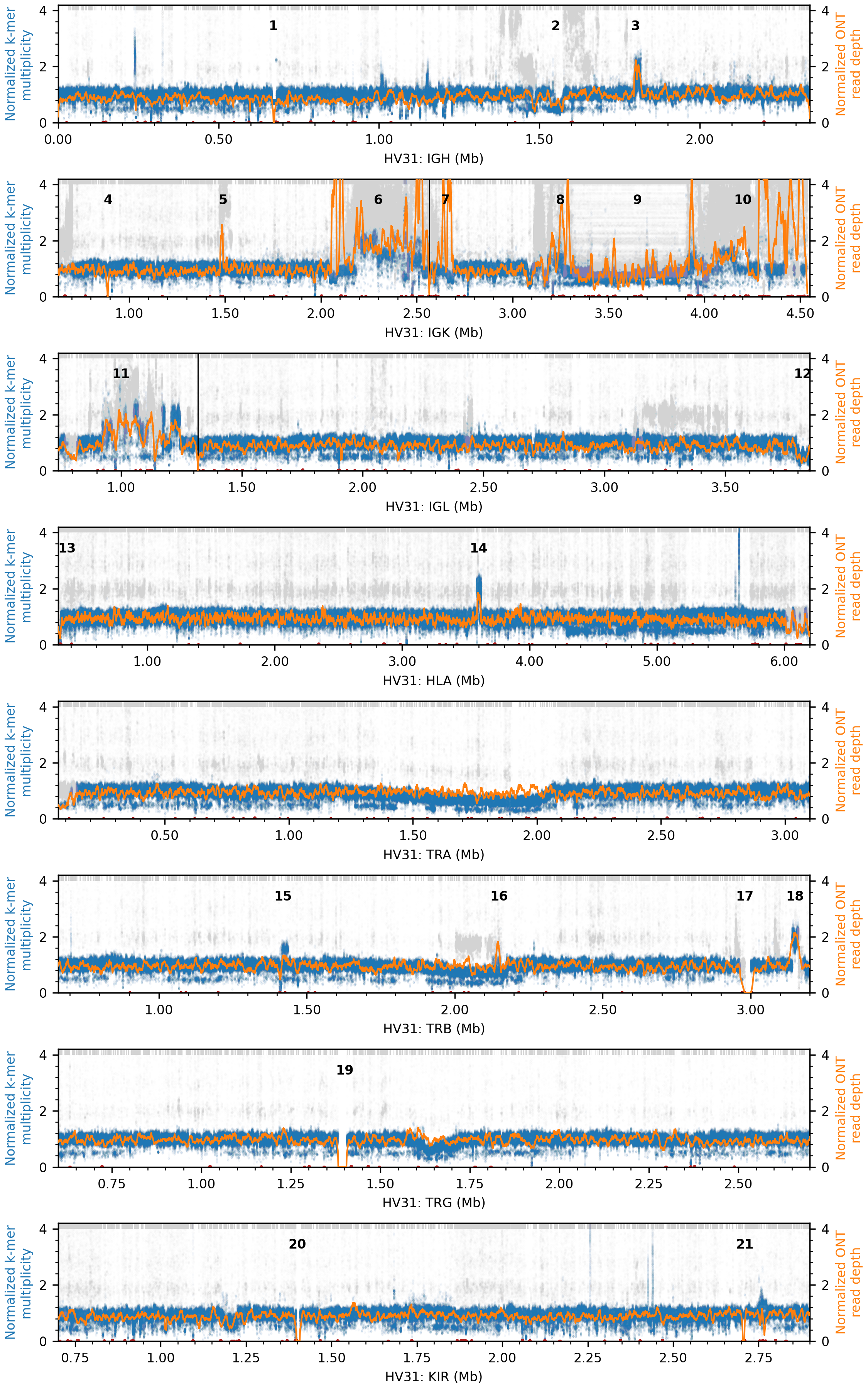
